# Characterisation of TbSmee1 indicates that endocytosis is required for access of surface-bound cargo to the trypanosome flagellar pocket

**DOI:** 10.1101/2022.03.15.484455

**Authors:** Daja Schichler, Antonia Konle, Eva-Maria Spath, Sina Riegler, Alexandra Klein, Anna Seleznev, Sisco Jung, Timothy Wuppermann, Noah Wetterich, Alyssa Borges, Elisabeth Meyer-Natus, Katharina Havlicek, Sonia Pérez Cabrera, Korbinian Niedermüller, Sara Sajko, Maximilian Dohn, Xenia Malzer, Emily Riemer, Tuguldur Tumurbaatar, Kristina Djinovic-Carugo, Gang Dong, Christian J. Janzen, Brooke Morriswood

## Abstract

All endo- and exocytosis in the African trypanosome *Trypanosoma brucei* occurs at a single subdomain of the plasma membrane. This subdomain, the flagellar pocket, is a small vase-shaped invagination containing the root of the cell’s single flagellum. Several cytoskeleton-associated multiprotein complexes are coiled around the neck of the flagellar pocket on its cytoplasmic face. One of these, the hook complex, was proposed to affect macromolecule entry into the flagellar pocket lumen. In previous work, knockdown of the hook complex component TbMORN1 resulted in larger cargo being unable to enter the flagellar pocket. In this study, the hook complex component TbSmee1 was characterised in bloodstream form *Trypanosoma brucei* and was found to be essential for cell viability. TbSmee1 knockdown resulted in flagellar pocket enlargement and impaired access to the flagellar pocket membrane by surface-bound cargo, similar to depletion of TbMORN1. Unexpectedly, inhibition of endocytosis by knockdown of clathrin phenocopied TbSmee1 knockdown, suggesting that endocytic activity itself is a prerequisite for the entry of surface-bound cargo into the flagellar pocket.

**Summary:** Characterisation of the essential trypanosome protein TbSmee1 suggests that endocytosis is required for flagellar pocket access of surface-bound cargo.

## Introduction

The flagellated parasitic protist *Trypanosoma brucei* lives in the bloodstream of its mammalian hosts, in continuous exposure to the immune system. Endo- and exocytosis are therefore critical to the parasite’s survival, as they are used to remove bound antibodies from the cell surface and scavenge macromolecular nutrients from the surroundings (Borst and Fairlamb, 1998; Engstler et al., 2007; Webster et al., 1990). Remarkably, all endo- and exocytosis in *T. brucei* occurs at just a single subdomain of the plasma membrane, despite the overall rate of endocytosis being comparable to a professional phagocyte (Engstler et al., 2004; Grunfelder et al., 2003). This subdomain is a small vase-shaped invagination that houses the root of the cell’s single flagellum, and is called the flagellar pocket (Halliday et al., 2021; Lacomble et al., 2009).

Molecules that enter the flagellar pocket, either in the fluid phase or when bound to the surface of the parasite (here “surface-bound”), are rapidly internalised by clathrin-mediated endocytosis, and then routed to the endosomal/lysosomal system (Link et al., 2021; Overath and Engstler, 2004). Depletion of clathrin results in a gross enlargement of the flagellar pocket due to an imbalance between endocytosis and exocytosis, and causes cargo accumulation inside the flagellar pocket (Allen et al., 2003). Although endocytosis of fluid phase and surface-bound cargo is relatively well-characterised in *T. brucei*, the exact mechanism(s) by which cargo initially enters the flagellar pocket are less well-understood.

Coiled around the neck of the flagellar pocket on its cytoplasmic face are a number of poorly-characterised cytoskeleton-associated complexes, which might contribute to these processes (Esson et al., 2012; Halliday et al., 2019)(Fig. 1A). The flagellar pocket collar is a multiprotein complex shaped like a cuff bracelet, and demarcates the boundary between the flagellar pocket and the flagellar pocket neck. It contains the protein TbBILBO1, and a number of its other components such as FPC4 and BILBO2 have recently been characterised (Albisetti et al., 2017; Bonhivers et al., 2008; Florimond et al., 2015; Isch et al., 2021). The centrin arm is a centrin-containing kinked rod which appears to be involved in the biogenesis of various cytoskeleton-associated structures (Selvapandiyan et al., 2007; Shi et al., 2008). The hook complex is a hook-shaped multiprotein complex that sits atop the flagellar pocket collar and alongside the centrin arm. Characterised components include the proteins TbMORN1, BOH1, BOH2, and Bhalin (Broster Reix et al., 2021; Morriswood et al., 2009; Pham et al., 2020; Pham et al., 2019). Previous characterisation of TbMORN1 revealed that it is essential for the viability of bloodstream form *T. brucei* cells (Morriswood and Schmidt, 2015). Cells depleted of TbMORN1 have enlarged flagellar pockets, indicative of an endocytosis defect. In addition, although small endocytic reporters such as 10 kDa dextran (hydrodynamic radius ∼1.86 nm) were still capable of entering the enlarged flagellar pocket, the access of larger macromolecules such as the lectin concanavalin A (ConA, hydrodynamic radius ∼4.2 nm) or bovine serum albumin (BSA, hydrodynamic radius ∼ 3.51 nm) conjugated to 5 nm gold particles was blocked (Ahmad et al., 2007; Armstrong et al., 2004; Morriswood and Schmidt, 2015). This phenotype was also observed upon depletion of the hook complex protein Bhalin (Broster Reix et al., 2021). On this basis, it was proposed that the hook complex might be regulating the passage of large macromolecules through the flagellar pocket neck, and thereby mediating cargo entry into the flagellar pocket (Morriswood and Schmidt, 2015).

**Figure 1.**
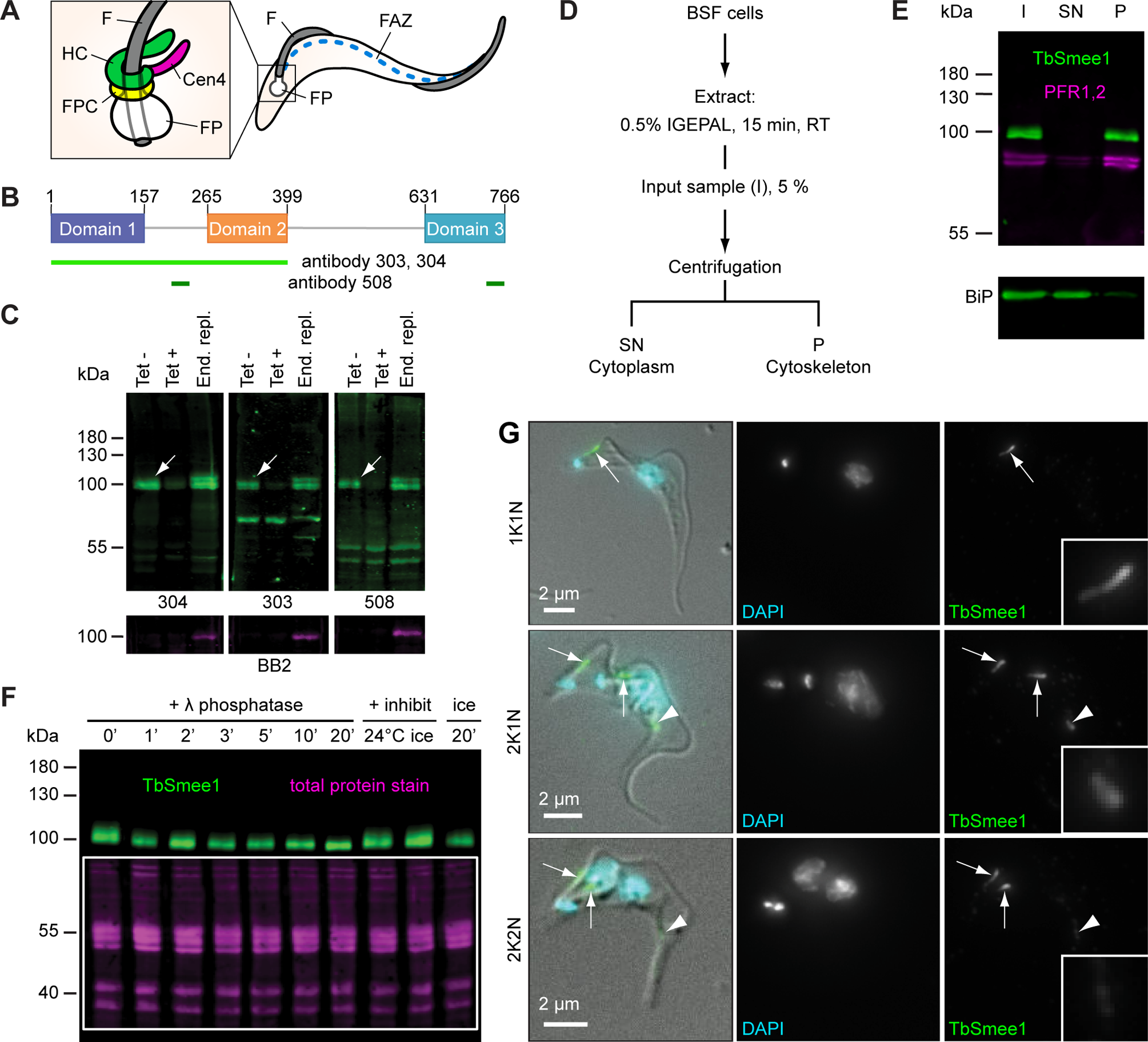
TbSmee1 is a cytoskeleton-associated phosphoprotein. (A) Schematic representation of a trypanosome cell with flagellar pocket (FP), flagellum (F), and flagellum attachment zone (FAZ) indicated. Inset shows the cytoskeleton-associated structures in the flagellar pocket neck area: hook complex (HC); flagellar pocket collar (FPC); centrin arm (Cen4). (B) Schematic representation of TbSmee1 (766 amino acids long) with the three predicted domains shown in purple, orange, and teal. Approximate amino acid ranges of each domain are indicated. The regions used as antigens for the generation of the three anti-TbSmee1 antibodies (303, 304, 508) are shown in light/dark green bars below the schematic. (C) Validation of the specificity of the three anti-TbSmee1 antibodies. Immunoblots against whole cell lysates from uninduced TbSmee1 RNAi cells (Tet −), induced TbSmee1 RNAi cells (Tet +), and Ty1-TbSmee1 endogenous replacement cells (End. repl.) using the three anti-TbSmee1 antibodies (304, 303, 508) are shown. All three antibodies detected an ∼ 85 kDa protein in the Tet - samples (upper panels, white arrows). The protein was strongly depleted after induction of TbSmee1 RNAi (Tet +). A second heavier band was detected in the endogenous replacement cells. This heavier band could also be detected using anti-Ty1 (BB2) antibodies (lower panels). Exemplary results from multiple (n>3) independent experiments are shown. (D) Fractionation scheme. Bloodstream form (BSF) cells were extracted with the non-ionic detergent IGEPAL and separated into cytoplasm and cytoskeleton fractions by centrifugation. (E) TbSmee1 is associated with the cytoskeleton. Immunoblot of whole-cell input (I), cytoplasmic supernatant (SN) and cytoskeletal pellet (P) fractions with anti-TbSmee1 antibodies. PFR1,2 (upper panel) and BiP (lower panel) were used as markers for the cytoskeleton and cytoplasm respectively. Equal fractions were loaded in each lane. Exemplary results from multiple (n>3) experiments are shown. (F) TbSmee1 is phosphorylated in vivo. Trypanosome cell lysates were incubated with λ phosphatase for the indicated times (0-20 min), followed by TbSmee1 detection by immunoblotting. The fuzzy appearance of the TbSmee1 was gradually lost over the timecourse. This was not observed in the presence of phosphatase inhibitors (+ inhibit), both at 24 °C and on ice, or when samples were kept on ice for 20 min. Equal loading of samples was confirmed using total protein stain (magenta). Exemplary results from multiple (n>3) independent experiments are shown. (G) TbSmee1 localisation through the cell cycle. Bloodstream form trypanosomes were extracted using non-ionic detergent, fixed, and labelled with anti-TbSmee1 antibodies. DNA was stained using DAPI. In 1K1N cells, TbSmee1 localised to a bar-shaped structure (arrow) close to the kinetoplast. In 2K1N cells the structure had replicated (arrows). TbSmee1 was additionally present on a third structure farther along the cell body (arrowhead). In 2K2N cells the signal intensity of the third structure was much weaker. Exemplary images from multiple (n>3) independent experiments are shown.

In this study, the hook complex protein TbSmee1 was characterised in bloodstream form *T. brucei*. TbSmee1 (Tb927.10.8820) was initially identified in a proximity labelling screen using TbMORN1 as bait, and shown to localise to the shank part of the hook complex (Morriswood et al., 2013). It was subsequently characterised in procyclic form *T. brucei*, the life cycle stage found within the tsetse fly vector (Perry et al., 2018). In bloodstream form *T. brucei,* depletion of TbSmee1 phenocopied TbMORN1 and Bhalin, with ConA, BSA and surface-bound antibodies unable to access the enlarged flagellar pocket. Surprisingly, clathrin-depleted cells also exhibited this flagellar pocket access phenotype, suggesting that the observed effects following depletion of the hook complex proteins are due to impaired endocytosis, and therefore that endocytosis is required for the entry of surface-bound cargo into the flagellar pocket.

## Results

### TbSmee1 is a cytoskeleton-associated phosphoprotein

Bioinformatic analysis predicted that TbSmee1 is composed of three structured domains separated by linker regions (Fig. 1B). Specifically, a multiple sequence alignment of the TbSmee1 protein with orthologues from other trypanosome species indicated the presence of three blocks of highly-conserved sequence (Fig. S1). These blocks of conserved sequence also corresponded to regions of predicted secondary structure (Fig. S2). Three-dimensional structural prediction by AlphaFold suggested a close association between the first two folded domains, and showed a good match with the secondary structural predictions (https://alphafold.ebi.ac.uk/entry/Q38A92) (Jumper et al., 2021; Wheeler, 2021).

Two polyclonal antibodies (303, 304) were raised against a recombinant TbSmee1(1-400) truncation, and a third (508) was raised against two peptides (Fig. 1B, green lines). The specificity of the antibodies was confirmed using immunoblots of whole-cell lysates obtained from TbSmee1 RNAi cells and 3xTy1-TbSmee1 endogenous replacement cells. In situ tagging of the *SMEE1* gene in the 3xTy1-TbSmee1 cells was confirmed by PCR analysis of genomic DNA (Fig. S3A,B). All three antibodies detected a protein of ∼ 85 kDa in the immunoblots (Fig. 1C, white arrows). This protein was strongly depleted in the RNAi (Tet +) samples (Fig. 1C). An extra band corresponding to 3xTy1-TbSmee1 was seen in lysates from the endogenous replacement (End. repl.) cells (Fig. 1C). This extra band was also detected using anti-Ty1 (BB2) antibodies, confirming its identity as Ty1-TbSmee1 (Fig 1C, lower panels).

To investigate TbSmee1 association with the cytoskeleton, cells were fractionated using non-ionic detergent (Fig. 1D). The fractions were analysed by immunoblotting, using antibodies against PFR1,2 as markers for the cytoskeleton, and against the ER chaperone BiP as a marker for the cytoplasm. TbSmee1 co-fractionated with PFR1,2, confirming that it was cytoskeleton-associated (Fig. 1E). Interestingly, the fuzzy appearance of the TbSmee1 band observed in immunoblots of whole-cell lysates was reduced or absent in the fractionation blots (compare Fig. 1C and Fig. 1E). TbSmee1 is heavily phosphorylated in vivo, and is a substrate and potential binding partner of the mitotic kinase TbPLK (Benz and Urbaniak, 2019; McAllaster et al., 2015; Nett et al., 2009; Urbaniak et al., 2013). The existence of different phosphoforms of a protein is known to cause a fuzzy appearance of bands in gels, so it seemed possible that exposure of TbSmee1 to endogenous phosphatases during the extraction step was causing dephosphorylation.

To investigate whether the band collapse could be attributed to dephosphorylation of TbSmee1, extracted cytoskeletons were incubated with exogenous phosphatase at various timepoints prior to immunoblotting. Incubation with exogenous phosphatase resulted in a progressive collapse of the TbSmee1 band over a 20 min period (Fig. 1F). This band collapse was not seen when the samples were in the presence of phosphatase inhibitors either at RT or on ice, or kept on ice (Fig. 1F, last three lanes). The fuzzy appearance of TbSmee1 in immunoblots could therefore be attributed exclusively to phosphorylation.

### TbSmee1 localises to the shank part of the hook complex

The localisation of endogenous and tagged TbSmee1 protein was analysed in bloodstream form *T. brucei* cells using immunofluorescence microscopy (Fig. 1G). All three main stages of the cell division cycle (1K1N, 2K1N, 2K2N) were analysed. In 1K1N cells (i.e. those with a single kinetoplast, K, and nucleus, N), TbSmee1 localised to a single structure near the flagellum base, consistent with the position of the hook complex (Fig. 1G, arrow). Interestingly, in 2K1N and 2K2N cells there was a third subpopulation of TbSmee1 at varying distances along the cell body in addition to the replicated hook complex (Fig. 1G, arrowheads). This subpopulation was fainter in 2K2N cells than 2K1N ones. The same TbSmee1 distributions were seen using all three anti-TbSmee1 antibodies; anti-Ty1 antibody labelling of 3xTy1-TbSmee1 cells also produced the same labelling pattern (Fig. S3C). In the endogenous replacement cells, the anti-TbSmee1 and anti-Ty1 signals colocalised (Fig. S3D).

To confirm the localisation of TbSmee1 at the hook complex, colabelling experiments were carried out. As expected, TbSmee1 strongly overlapped with the shank part of the hook complex as labelled by TbMORN1 and TbLRRP1 (Fig. 2A, B). TbSmee1 also strongly overlapped with the hook complex protein Tb927.10.3010. This 133 kDa protein was one of the top hits in the BioID screen using TbMORN1 (Morriswood et al., 2013). Consistent with the convention started with TbSmee1, Tb927.10.3010 was named TbStarkey1 after another of Captain Hook’s pirates. Two anti-peptide antibodies were generated against TbStarkey1, and their specificity was confirmed by immunoblotting and immunofluorescence imaging of TbStarkey1 RNAi cells (Fig. S4A, B). Strong overlap was seen between TbSmee1 and TbStarkey1 on the shank part of the hook complex (Fig. S4C). As expected, TbSmee1 showed only a partial overlap with TbCentrin4, a marker of the centrin arm, and little overlap with TbBILBO1, a marker of the flagellar pocket collar (Fig. 2C, D).

**Figure 2.**
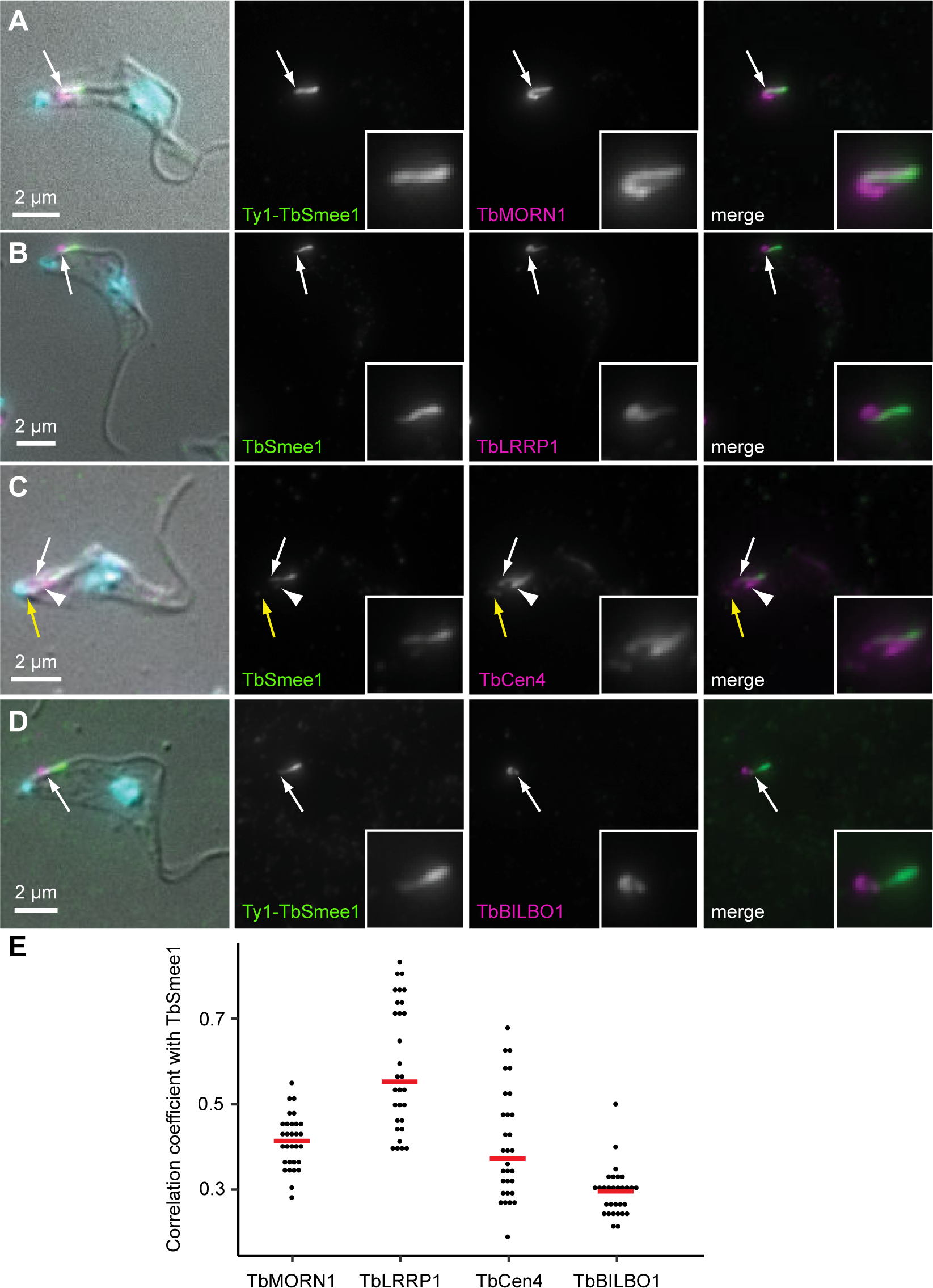
TbSmee1 is localised to the shank part of the hook complex. Bloodstream form trypanosomes were extracted with non-ionic detergent, fixed, and labelled with the indicated antibodies. Either wild-type or Ty1-TbSmee1 cells were used. Insets show an enlarged view of the hook complex region. (A) TbSmee1 overlaps with the shank part of the hook complex protein TbMORN1 (arrow). (B) TbSmee1 overlaps with the shank part of the hook complex protein TbLRRP1 (arrow). (C) TbSmee1 partially overlaps with TbCen4. TbCen4 is present at the basal and probasal bodies (yellow arrow), centrin arm (arrowhead), and a small additional projection (white arrow). (D) TbSmee1 does not overlap with the flagellar pocket collar protein TbBILBO1 (arrow). (E) Summary of measured correlation coefficients for each of the colabelling experiments; red bars show median values. TbSmee11 showed a moderate correlation with TbMORN1 (0.41) and TbLRRP1 (0.55) and a weak correlation with TbCen4 (0.37) TbBILBO1 (0.3). Each dot represents a single cell in the 1K1N stage (N = 30). All fluorescence images are maximum intensity z-projections, and an overlay with a single DIC section is shown. Overlap was manually confirmed in single z-slices. Results were obtained from multiple (n>3) independent experiments; exemplary images are shown.

To quantify the degree of colocalisation between TbSmee1 and the various hook complex and flagellar pocket collar marker proteins, correlation coefficients from the pairwise labelling experiments were calculated. Moderate correlation was seen between TbSmee1 and TbMORN1 and TbLRRP1, with lower values being measured for TbCentrin4 and TbBILBO1 (Fig. 2E).

### TbSmee1 is transiently associated with the flagellum attachment zone (FAZ) tip

The flagellum attachment zone (FAZ) is a cytoskeleton-associated apparatus that adheres the flagellum of *T. brucei* to the cell body (Sunter and Gull, 2016). During replication, a new FAZ is assembled and grows from its initiation point very close to the hook complex towards the anterior end of the cell (Sunter et al., 2015b; Zhou et al., 2015). It was previously shown that a tagged version of TbSmee1 is transiently localised to the tip of the new FAZ in replicating insect-stage *T. brucei* cells, in addition to its hook complex localisation (Perry et al., 2018). This dynamic localisation contrasts with other hook complex proteins such as TbMORN1, which are localised exclusively to the hook complex.

To investigate whether the endogenous TbSmee1 subpopulation seen in 2K1N and 2K2N BSF cells corresponded to the FAZ tip, TbSmee1 and the FAZ marker FAZ1 were analysed at different stages of the cell division cycle. As expected, the TbSmee1 present at the hook complex showed a partial overlap with the posterior end of the FAZ (Fig. 3A, arrow). The extra TbSmee1 subpopulation in 2K1N and 2K2N cells overlapped with the tip of the new FAZ (Fig. 3A, arrowheads). The TbSmee1 signal got fainter as the new FAZ elongated, and was quite faint in the 2K2N stage. TbSmee1 was sometimes faintly observed along the full length of the new FAZ in replicating cells, and upon careful observation, a TbSmee1 signal could also be detected at the tip of the old FAZ as well, even in 1K1N cells.

**Figure 3.**
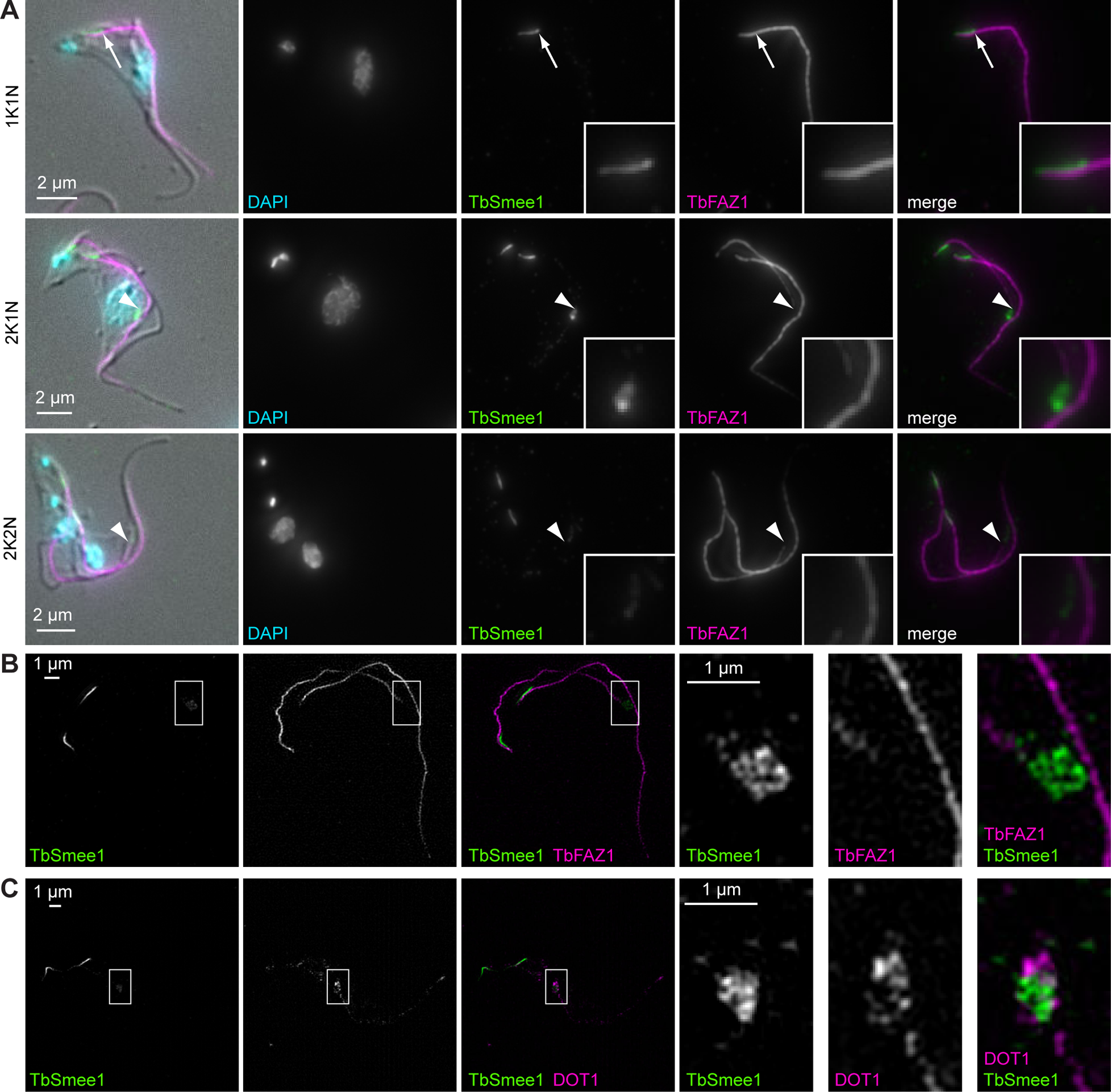
TbSmee1 is associated with the tip of the elongating new FAZ in replicating cells. (A) Bloodstream form trypanosomes were extracted with non-ionic detergent, fixed, and labelled with the indicated antibodies. DNA was stained with DAPI. Maximum intensity projections are shown, with a single DIC z-slice overlaid. In 1K1N cells, TbSmee1 partially overlapped (arrow) with the posterior end of the FAZ. In 2K1N and 2K2N cells, the additional TbSmee1 structure (arrowhead) was present at the tip of the elongating new FAZ. Exemplary images from multiple (n>3) independent experiments are shown. (B) SIM image of the same preparations. The TbSmee1 structure (inset) lay just ahead of the elongating FAZ tip. (C) SIM image of a detergent-extracted cell colabelled for TbSmee1 and DOT1. DOT1 appeared to envelop and partially overlap with TbSmee1.

Using structured illumination microscopy, TbSmee1 was observed to be slightly in front of the tip of the new FAZ filament (Fig. 3B). This suggested that TbSmee1 might be present at the “groove”. The groove is a structure involved in remodelling of the microtubule cytoskeleton during cell replication in bloodstream form *T. brucei* (Hughes et al., 2013; Smithson et al., 2022). It can be detected using the monoclonal antibody DOT1. TbSmee1 appeared to be enveloped by the DOT1 labelling, confirming its presence at the groove (Fig. 3C). In summary, in replicating cells TbSmee1 is present at the groove in addition to the hook complex, and travels in front of the newly-assembling FAZ filament.

### The C-terminal part of TbSmee1 is required for targeting to the hook complex

To determine what parts of TbSmee1 primary structure are responsible for targeting to the hook complex, a series of truncation constructs based on the predicted domain architecture of TbSmee1 were designed (Fig. 4A). Cell lines were generated that could inducibly express each of these truncations with an N-terminal Ty1 tag. The presence of the ectopic transgenes in the genomes of these cells was confirmed by PCR analysis of genomic DNA (Fig. S5A). To determine the localisation of each construct, and whether it could associate with the cytoskeleton, detergent-extracted cells were analysed using immunofluorescence microscopy. It should be noted that both endogenous *SMEE1* alleles were still present, meaning that targeting of the truncation constructs was being assayed in the presence of the endogenous Smee1 protein.

**Figure 4.**
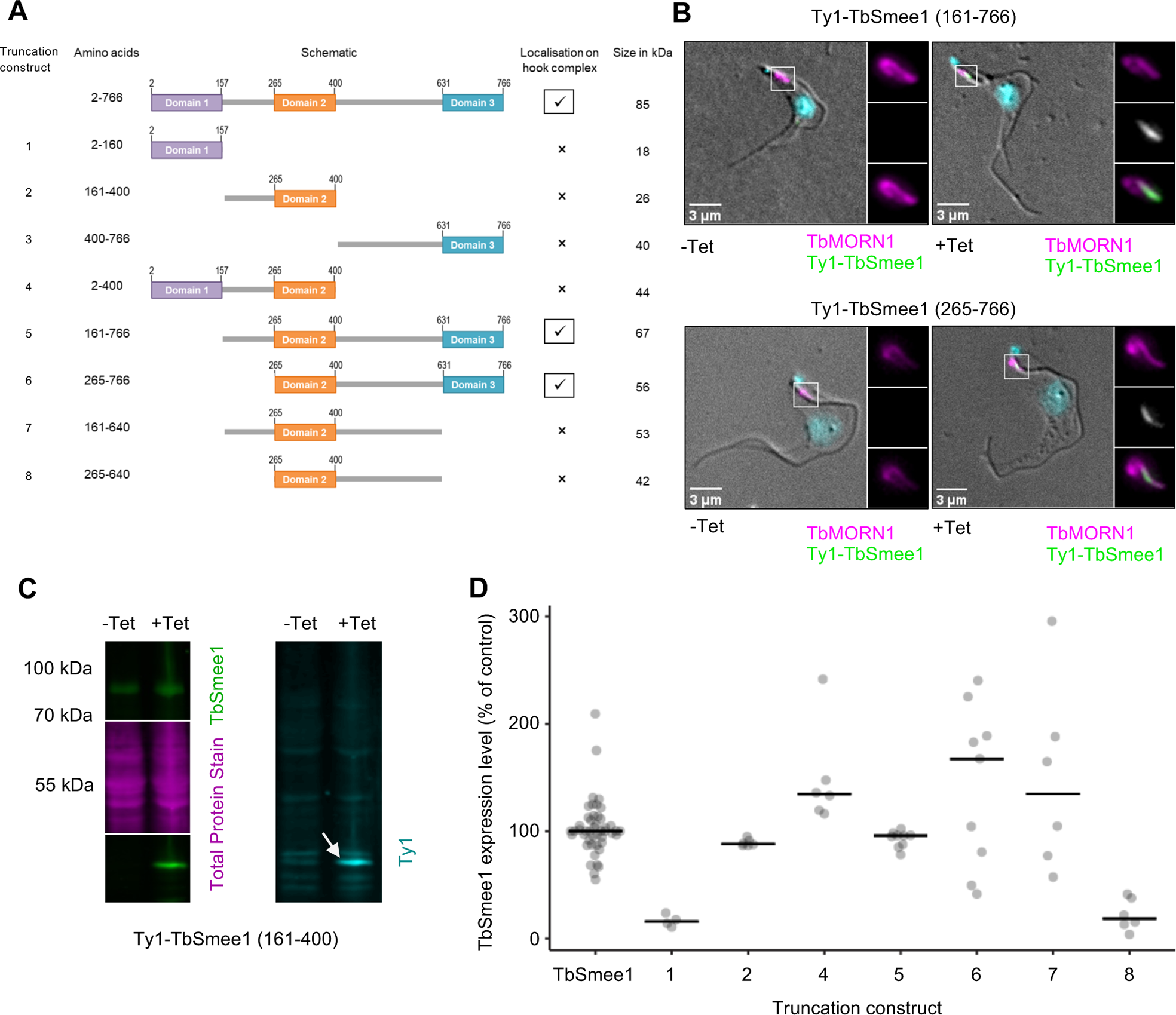
TbSmee1 domains 2 and 3 are required for targeting to the hook complex. (A) Schematics of the 8 TbSmee1 truncation constructs tested, along with details of amino acid ranges, localisation, and size in kDa. (B) TbSmee1 domains 2 and 3 are required for localisation to the hook complex. Stably-transfected cells inducibly expressing the indicated TbSmee1 truncations were used. Ty1-TbSmee1 was detected using anti-Ty1 antibodies. In the absence of induction (−Tet), no signal overlapping with endogenous TbMORN1 was seen. After induction of expression (+Tet), the Ty1-TbSmee1 truncations overlapped with the shank part of TbMORN1 (insets). Images shown are maximum intensity projections of the fluorescence channels overlaid with a single DIC z-slice. Exemplary images are shown. (C) Confirmation of Ty1-TbSmee1 truncation expression. Whole-cell lysates from control (−Tet) cells and cells expressing Ty1-TbSmee1 truncations (+Tet) were immunoblotted using anti-TbSmee1 (left panel) and anti-Ty1 (right panel) antibodies. Total protein stain (magenta) was used as a loading control. In the exemplary blot shown, the Ty1-TbSmee1 was detected at ∼ 44 kDa, as expected. (D) Quantification of immunoblotting data. Anti-TbSmee1 signals in the immunoblots were normalised relative to total protein staining. TbSmee1 levels in uninduced control cells (TbSmee1) were expressed relative to the mean value of all control samples for each clone in each experiment. Ty1-TbSmee1 levels were expressed relative to the levels of endogenous TbSmee1 for each clone in each experiment. The data shown were obtained from two independent experiments with each Ty1-TbSmee1 truncation; each experiment used three separate clones.

TbSmee1(161-766) correctly localised to the hook complex, indicating that Domain1 is not necessary for targeting (Fig. 4B). Of note, no dominant negative effects on cell growth were observed upon overexpression of the TbSmee1(161-766) construct. TbSmee1(265-766) also localised correctly, indicating that the predicted linker region between Domain1 and Domain2 is not necessary for localisation (Fig. 4B). No other truncations were observed to localise to the hook complex, making TbSmee1(265-766) the smallest construct to localise correctly. Expression of all TbSmee1 truncations was confirmed by immunoblotting with anti-TbSmee1 and anti-Ty1 antibodies (Figs 4C, S5A). Quantification of the immunoblots indicated that most were expressed at around the level of the endogenous protein (Fig. 4D).

Interestingly, TbSmee1(161-766) also localised to the FAZ tip in replicating cells (Fig. S5B). TbSmee1(265-766) did not localise to the FAZ tip, suggesting that the linker region between Domain1 and Domain2 (aa 161-264) might be required for targeting to this structure. In support of this hypothesis, TbSmee1(2-400) was found to localise to the FAZ tip despite not localising to the hook complex (Fig. S5C).

### TbSmee1 is essential for the viability of BSF *T. brucei*, and its depletion causes gross enlargement of the flagellar pocket

The effects of TbSmee1 depletion were analysed using tetracycline-inducible RNAi. Induction of RNAi (Tet+) resulted in a rapid cessation of population growth after around 24 h (Fig. 5A). Visual inspection of the stalled populations at 48 h and 72 h post-induction showed widespread lysis, confirming that TbSmee1 is essential for the viability of bloodstream form cells in vitro. A shorter timecourse with higher sampling frequency indicated that the growth defect began after around 20 h of RNAi, and was already clear at 24 h (Fig. 5A, inset).

**Figure 5.**
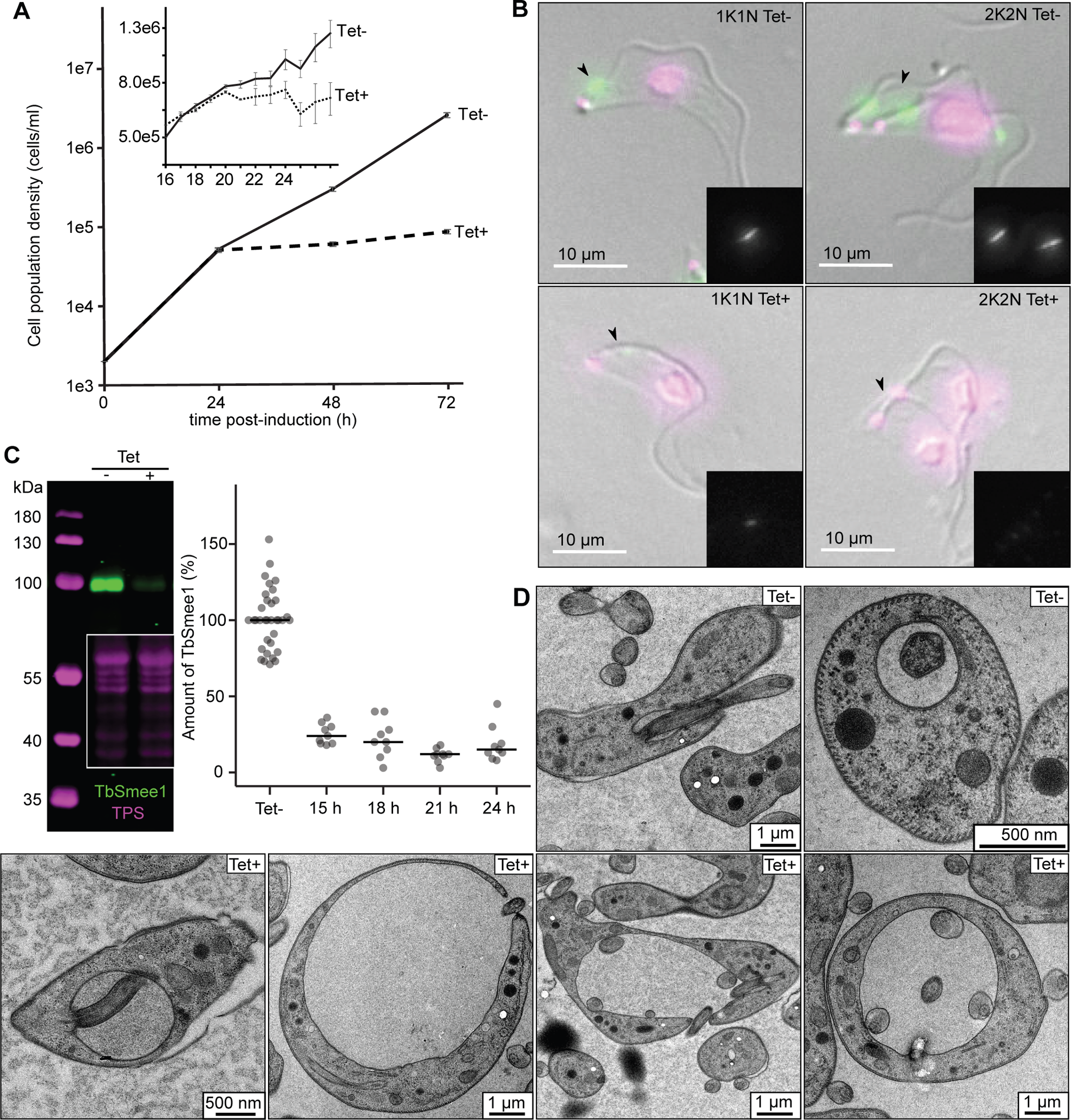
TbSmee1 is essential for the viability of bloodstream form *T. brucei*. (A) Depletion of TbSmee1 causes a strong growth defect. The population density of uninduced control (Tet-) and TbSmee1-depleted (Tet+) cells were measured at regular intervals after induction of RNAi over a 72 h timecourse. The insert shows data from experiments with readings taken at hourly intervals. A growth defect was evident from 21 h post-induction onwards. (B) Confirmation of TbSmee1 depletion at the single-cell level. Uninduced control (Tet-) and TbSmee1-depleted (Tet+) RNAi cells were extracted, fixed, and labelled with anti-TbSmee1 antibodies (green). DNA was labelled with DAPI (magenta). Maximum intensity projections of the fluorescence channels are shown overlaid with a single DIC z-slice. Insets show the TbSmee1 signal. Exemplary 1K1N and 2K2N cells from each condition are shown. (C) Confirmation and quantification of TbSmee1 depletion. Whole-cell lysates from control (Tet-) and TbSmee1-depleted (Tet+) RNAi cells were immunoblotted using antibodies against TbSmee1 (green). Total protein stain (TPS, magenta) was used for signal normalisation. An exemplary immunoblot is shown. Normalised TbSmee1 signals in +Tet cells were expressed relative to the −Tet signal for each clone in each experiment. TbSmee1 −Tet signals for each clone in each experiment were expressed relative to the mean of all TbSmee1 −Tet values in the dataset. TbSmee1 depletion was quantified at the indicated timepoints post-induction. (D) Depletion of TbSmee1 results in an enlargement of the flagellar pocket. Electron microscopy images of control (Tet-) and TbSmee1-depleted (Tet+) RNAi cells are shown. The cells were fixed 24 h post-induction. All data in the experiments shown in this figure were obtained from multiple (n≥3) independent experiments, each using three separate clones.

TbSmee1 depletion was confirmed at the single-cell level using immunofluorescence microscopy (Fig. 5B). TbSmee1 signal was lost from both the hook complex and FAZ tip. The kinetics of TbSmee1 protein depletion on either side of the onset of the growth defect were assessed by immunoblotting. TbSmee1 protein levels were reduced to ∼ 20-25% at 15 h and 18 h post-induction, with a further reduction to around 10-15% from 21 h onwards (Fig. 5C).

The effect of TbSmee1 knockdown on cell cycle progression was assessed by quantifying the numbers of 1K1N, 2K1N, 2K2N, and abnormal cell types in the same time window (Fig. S6A). An increase in 2K1N and a decrease in 1K1N cells was visible from 21 h post-induction onwards, followed by an increase in 2K2N cells at the 24 h timepoint. This indicated that cell cycle progression was inhibited from around 21 h after induction of RNAi, correlating with the onset of the growth defect. To summarise, TbSmee1 protein was already significantly depleted at 15 h, with subsequent effects on population growth and cell cycle progression visible from around 21 h onwards.

The effects of TbSmee1 depletion on a panel of hook complex proteins and components of the centrin arm and flagellar pocket collar were systematically evaluated (Fig. S7A-E). TbSmee1 depletion for 24 h did not result in observable effects on the expression levels or localisation of any of the candidates. No flagellum detachment was observed either. Conversely, depletion of TbMORN1 for just 16 h resulted in reductions in the levels of TbSmee1 and TbStarkey1 (Fig. S7B, F-H). The lack of observable structural changes to the hook complex, flagellar pocket collar, and centrin arm following TbSmee1 depletion implied that the lethal phenotype might be due to a loss of a specific cellular function, rather than destabilisation of structural complexes.

To determine the ultrastructural changes caused by TbSmee1 depletion, the cells were imaged by electron microscopy after high-pressure freezing. High-pressure freezing was used, as this approach gives better morphological preservation than chemically fixing the cells in the growth media. The most obvious morphological effect of TbSmee1 depletion was the gradual accumulation of cells with enlarged flagellar pockets (Fig. 5D). This presumably was the cause of the lethality phenotype, as depletion of TbMORN1 was also shown to result in progressive enlargement of the flagellar pocket until cells rounded up and lysed (Morriswood and Schmidt, 2015). Interestingly however, depletion of TbStarkey1 also frequently resulted in the generation of cells with enlarged flagellar pockets, despite causing no growth defect (Fig. S6B-D). This, together with the recently-published characterisation of Bhalin (Broster Reix et al., 2021), means that depletion of four separate hook complex proteins - TbMORN1, TbSmee1, TbStarkey1, Bhalin - results in flagellar pocket enlargement, though with varying magnitudes of effect.

### Flagellar pocket enlargement is an early consequence of TbSmee1 depletion

Although flagellar pocket enlargement appears to be a consistent phenotype resulting from hook complex protein depletion, it is not an uncommon phenotype in bloodstream form RNAi cells. Importantly, it can be either a direct or an indirect consequence of protein depletion (Ali et al., 2014; Allen et al., 2003; Hall et al., 2004; Hall et al., 2005; Price et al., 2007).

To determine whether flagellar pocket enlargement was an early/direct consequence of TbSmee1 depletion (i.e. occurring very soon after or even before the onset of the growth defect), a fluorescent, fixable 10 kDa dextran reporter was used. Dextran is a polysaccharide that traffics in the fluid phase and is well-established as an endocytic marker. Control (− Tet) and TbSmee1-depleted (+ Tet) cells were incubated on ice in order to block endocytosis (Brickman et al., 1995). The cells were then incubated with the labelled dextran for 15 min to allow it to enter the flagellar pocket, and fixed afterwards (Fig. 6A). The magnitude of the dextran signal should therefore be proportional to flagellar pocket volume. Visual analysis of the cells confirmed that dextran visibly labelled the flagellar pocket, with a much greater signal seen for cells with an enlarged flagellar pocket (Fig. 6B).

**Figure 6.**
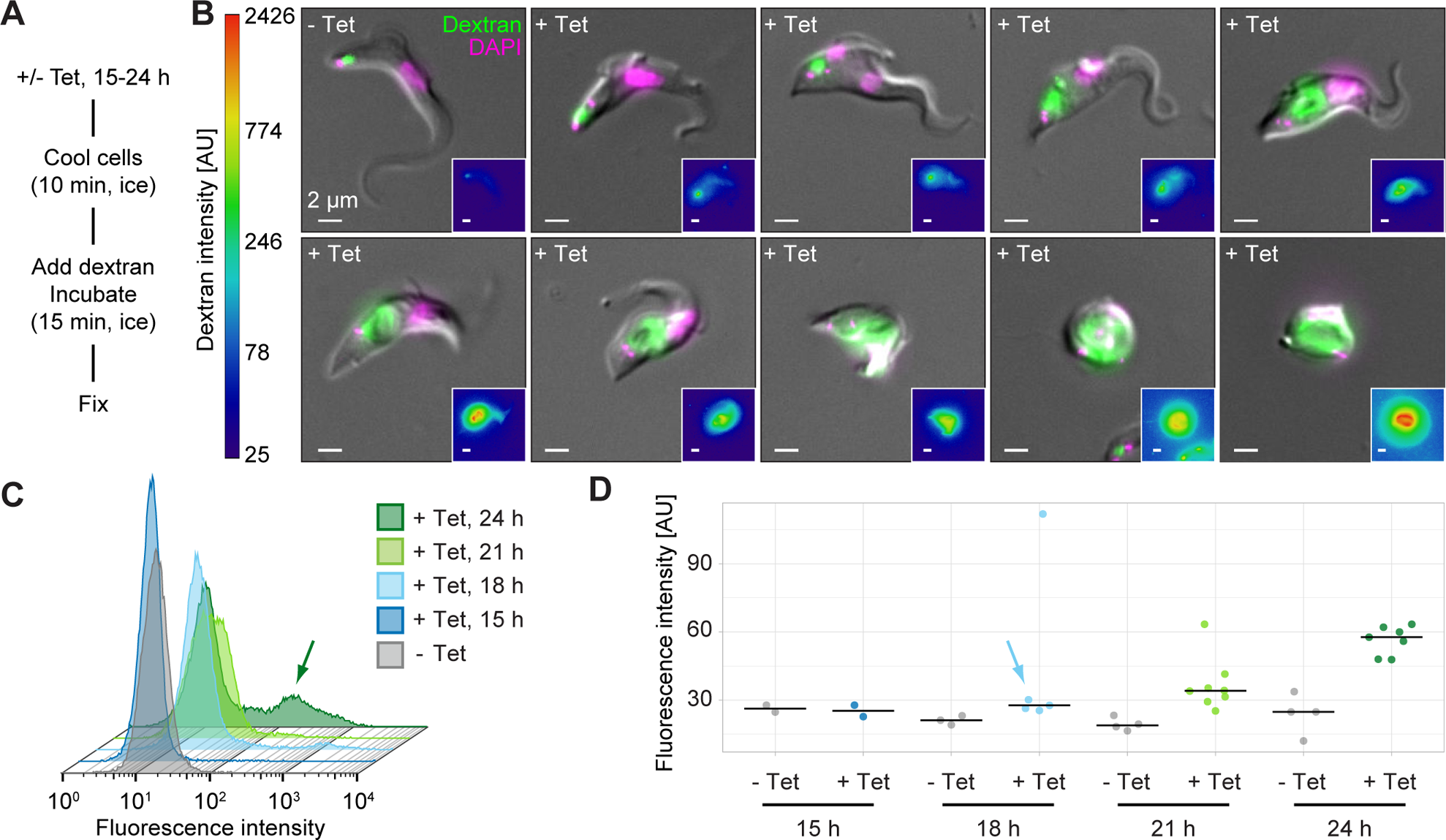
Flagellar pocket enlargement is an early consequence of TbSmee1 depletion. (A) Summary of dextran uptake protocol. The use of temperature blocks inhibits endocytosis, allowing dextran to enter the flagellar pocket but not be internalised. (B) The intensity of the dextran signal reports on flagellar pocket volume. Cells were incubated with fluorophore-conjugated dextran (green) prior to being fixed and imaged using fluorescence microscopy. DNA was labelled with DAPI (magenta). Exemplary cells from control (− Tet) and TbSmee1-depleted (+ Tet) conditions are shown. The + Tet cells (18, 21, or 24 h of induction) exhibited various degrees of flagellar pocket enlargement and progressive morphological aberration. Insets show the dextran signal with a log-scale LUT. (C) Flow cytometry analysis of control (− Tet) and TbSmee1-depleted (+ Tet) cells incubated with fluorescent dextran at various timepoints after induction of RNAi. At later timepoints there is a clear emergence of a subpopulation of cells with much greater fluorescence intensity (arrow). Exemplary traces from a single experiment are shown. (D) Quantification of flow cytometry data. The geometric mean of the fluorescence intensity in control (− Tet) and TbSmee1-depleted (+ Tet) cells was measured at various timepoints after induction of RNAi. Bars indicate median values. The population mean of the + Tet cells was clearly higher at the later timepoints; a visible shift was visible as early as 18 h post-induction (arrow). All data in the experiments shown in this figure were obtained from multiple (n≥2) independent experiments; each experiment used three separate clones.

Flow cytometry was then used for high-throughput, unbiased, and quantitative analysis of the cells (Fig. 6C). At 15 h post-induction, no difference between TbSmee1-depleted cells and controls was observed. At 18 h and 21 h post-induction a slight “shoulder” on the +Tet traces became visible, indicating the emergence of cells with higher fluorescence values than seen in controls. By 24 h post-induction there was a clear hump visible in the traces, indicating a subpopulation with fluorescence values sometimes two orders of magnitude greater than those seen in control cells (Fig. 6C, arrow). Quantification of the flow cytometry data from multiple experiments showed that average fluorescence intensity of the whole + Tet population was noticeably higher than controls at 21 h and 24 h post-induction (Fig. 6D). Even at 18 h post-induction, i.e. before the onset of the growth defect, there was already a clear increase in the average fluorescence intensity in the + Tet population (Fig. 6D, blue arrow). This strongly suggests that flagellar pocket enlargement is an early and probably direct consequence of TbSmee1 depletion.

### TbSmee1 depletion results in impaired flagellar pocket access of surface-bound cargo

It was previously shown that the ability of large cargo to enter the flagellar pocket is affected after knockdown of the hook complex protein TbMORN1 (Morriswood and Schmidt, 2015). Specifically, the fluid phase marker 10 kDa dextran accumulates in the enlarged flagellar pocket of TbMORN1-depleted cells, while larger fluid phase cargo such as BSA-5 nm gold and large surface-bound cargo such as ConA (which binds to surface glycoproteins) do not access the flagellar pocket lumen.

To test whether TbSmee1 knockdown also impairs flagellar pocket access, TbSmee1 RNAi cells were incubated with both dextran and ConA to simultaneously monitor the uptake of fluid phase and surface-bound cargo. The cells were first incubated on ice to block endocytosis, and then with the reporters (also on ice) to allow ConA to bind and dextran to enter the flagellar pocket. The cells were then shifted to 37°C to reactivate endocytosis, and subsequently fixed and imaged (Fig. 7A).

**Figure 7.**
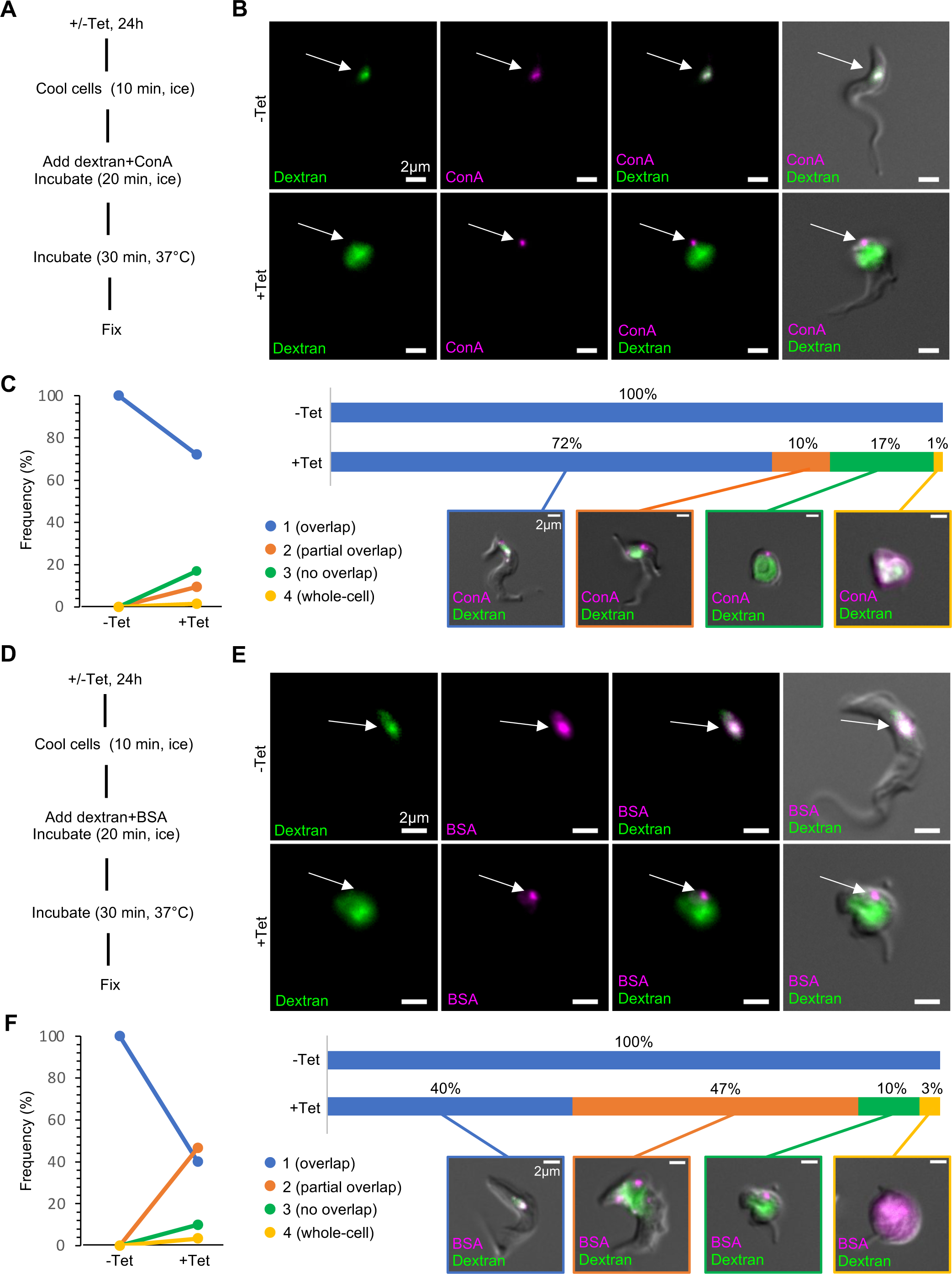
Surface-bound reporters cannot enter the flagellar pocket of TbSmee1-depleted cells. (A) Summary of paired dextran+ConA uptake protocol. (B) ConA is unable to enter the enlarged flagellar pocket of TbSmee1-depleted cells. In control (−Tet) cells, the dextran and ConA signals strongly overlapped (arrow, upper panels). In TbSmee1-depleted cells (+Tet), little to no overlap between the two reporters was observed (arrow, lower panels). Maximum intensity projections of the fluorescence channels are shown overlaid with a single DIC z-slice. (C) Quantification of dextran+ConA uptake experiments. Cells from control (−Tet) and TbSmee1-depleted (+Tet) samples were manually classified into four categories (1-4) based on the degree of overlap between the ConA and dextran reporters. Results are shown as a slope chart (left) and stacked bar chart (right); exemplary cells for each category are shown. Percentages represent total cell counts (442 cells) that were obtained from multiple independent experiments (n>3); each experiment included three separate clones. (D) Summary of paired dextran+BSA uptake protocol. (E) BSA is unable to enter the enlarged flagellar pocket of TbSmee1-depleted cells. (F) Quantification of dextran+BSA uptake experiments. Percentages represent total cell counts (103 cells) that were obtained from multiple independent experiments (n>3); each experiment included three separate clones.

In control (− Tet) cells, both dextran and ConA strongly overlapped in the part of the cell corresponding to the endosomal/lysosomal system (Fig. 7B, −Tet, arrow). To confirm that the internalised material was being trafficked to the lysosome, the cells were labelled with antibodies specific for the lysosomal enzyme p67 (Kelley et al., 1999). The dextran reporter was not compatible with immunolabelling, but the ConA signal clearly overlapped with the lysosome marker p67 (Fig. S8, −Tet cells). In TbSmee1-depleted (+ Tet) cells, dextran filled the enlarged flagellar pocket while ConA was restricted to one or two small foci that appeared to be on the cell surface (Fig. 7B, +Tet, arrow). No overlap was seen between the dextran and ConA labels, suggesting that ConA was not able to enter the flagellar pocket. In addition, no overlap was observed between ConA and the lysosome marker p67 (Fig. S8, +Tet cells).

To quantify these observations, the cells were grouped into 4 categories: 1, complete overlap between the two labels; 2, partial overlap between the two labels; 3, no overlap between the two labels; 4, whole-cell labelling. Whole-cell labelling occurs when the cell has lost integrity, and labelling is found throughout the cytoplasm. Control (−Tet) cells all showed complete overlap between the dextran and ConA, while cells depleted of TbSmee1 for 24 h showed >25% of cells with partial or no overlap between the cargoes (Fig. 7C).

To further validate these observations, the experiments were repeated using fluorescently-labelled BSA (Fig. 7D). BSA traffics in the fluid phase and is a physiological cargo, unlike ConA (Coppens et al., 1987). As expected, in control cells there was strong overlap between dextran and BSA from the endosomal/lysosomal system (Fig. 7E, −Tet, arrow). In TbSmee1-depleted cells, there was once again no overlap between the reporters, indicating that BSA was unable to access the enlarged flagellar pocket. Surprisingly, despite BSA being fluid-phase cargo there was a punctate signal analogous to that seen with ConA, suggesting that it could at least partly bind to the cell surface (Fig. 7E, +Tet, arrow). Quantification of the data showed that the effect on BSA was more pronounced than that seen for ConA, with >55% of TbSmee1-depleted cells showing no or only partial overlap between the two reporters after 24 h of RNAi (Fig. 7F).

### Endocytosis is required for flagellar pocket access of surface-bound cargo

The inability of surface-bound cargo to enter the flagellar pocket of TbMORN1- and TbSmee1-depleted cells could either be due to a defect in endocytosis (indicated by the enlargement of the flagellar pocket), or a secondary effect.

To distinguish between these possibilities, the assays were repeated following clathrin depletion (Fig. 8). Clathrin is an essential endocytic coat protein and its loss results in a block in endocytosis. This results in the expected enlargement of the flagellar pocket and ultimately cell lysis, and in fact this phenotype was first described as a result of clathrin depletion (Allen et al., 2003). Studies of clathrin in trypanosomes have however focused on whether or not cargo is endocytosed, but not whether or not it can enter the flagellar pocket prior to encountering the block in internalisation. In control cells, as expected, there was strong overlap between the dextran and ConA cargoes in the region of the endolysosomal system (Fig. 8B, −Tet, arrow). In clathrin-depleted cells however, there was again no overlap observed between the two reporters, and ConA did not appear able to access the enlarged flagellar pocket (Fig. 8B, +Tet, arrow). Quantification of the data showed that >50% of clathrin-depleted cells showed either partial or no overlap between the two reporters after 19 h of RNAi (Fig. 8C). The same effect was seen in assays using dextran and BSA, with little to no overlap between the reporters in clathrin-depleted cells (Fig. 8D-F). These results indicated that inhibition of endocytosis by itself resulted in the failure of either ConA or BSA to enter the enlarged flagellar pocket.

**Figure 8.**
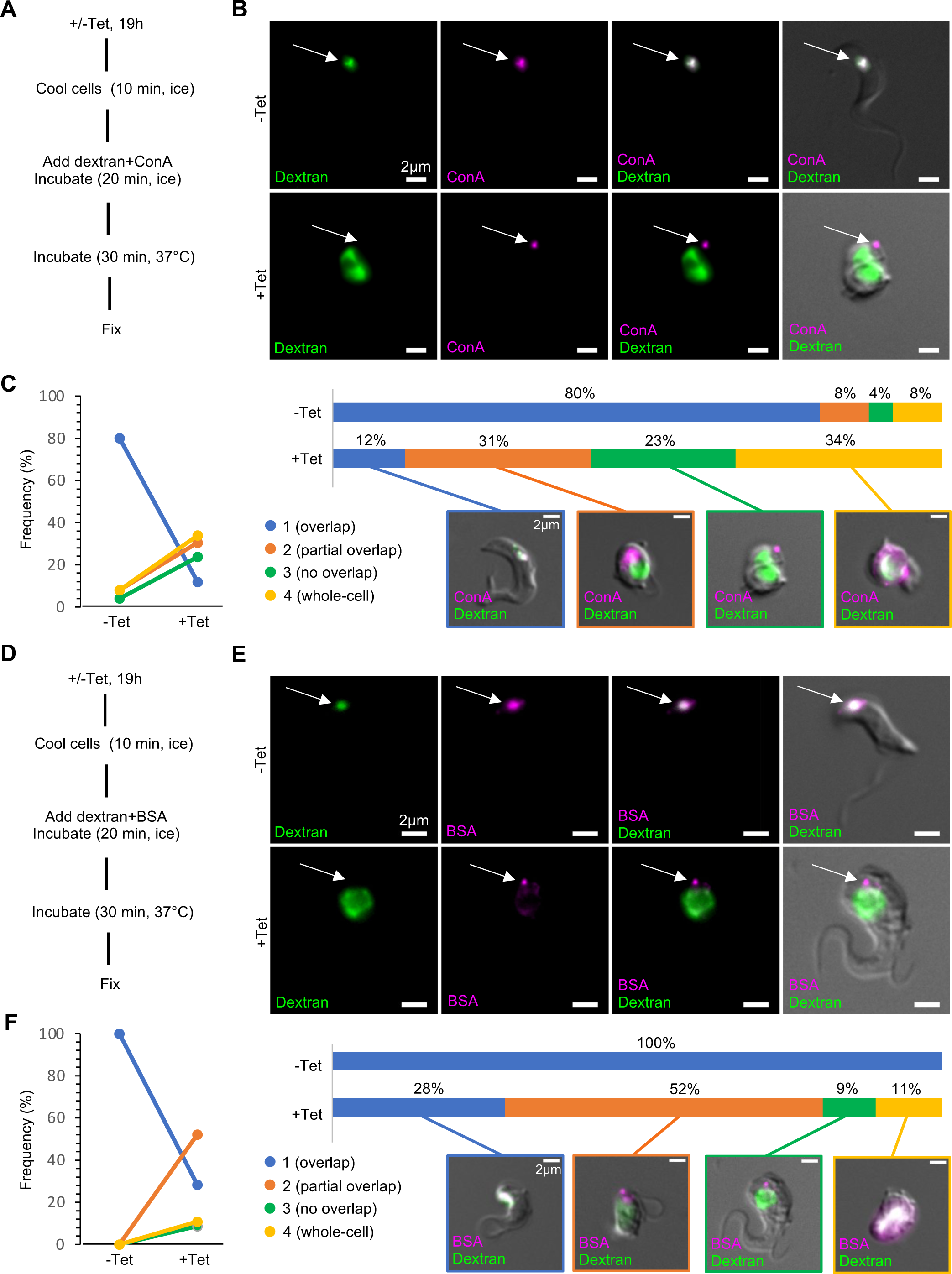
Blockade of surface reporters is due to an inhibition of endocytosis. (A) Summary of paired dextran+ConA uptake protocol. (B) ConA is unable to enter the enlarged flagellar pocket of clathrin-depleted cells. In control (−Tet) cells, the dextran and ConA signals strongly overlapped (arrow, upper panels). In clathrin-depleted cells (+Tet), little to no overlap between the two reporters was observed (arrow, lower-panels). Maximum intensity projections of the fluorescence channels are shown overlaid with a single DIC z-slice. (C) Quantification of dextran+ConA uptake experiments. Cells from control (−Tet) and clathrin-depleted (+Tet) samples were manually classified into four categories (1-4) based on the degree of overlap between the ConA and dextran reporters. Results are shown as a slope chart (left) and stacked bar chart (right); exemplary cells for each category are shown. Percentages represent total cell counts (84 cells) that were obtained from multiple independent experiments (n>3); each experiment included three separate clones. (D) Summary of paired dextran+BSA uptake protocol. (E) BSA is unable to enter the enlarged flagellar pocket of clathrin-depleted cells. (F) Quantification of dextran+BSA uptake experiments. Percentages represent total cell counts (58 cells) that were obtained from multiple independent experiments (n>3); each experiment included three separate clones.

### Endocytosis is required for flagellar pocket access of surface-bound antibodies

One of the main functions of endocytosis in trypanosomes is to remove any antibodies bound to the surface glycoprotein coat. To determine whether the effects observed above using ConA and BSA reporters also apply to antibodies, another set of uptake assays were carried out. As the uptake of surface-bound antibodies occurs in a matter of seconds, a slightly modified assay protocol was used (Fig. 9A). The cells were first incubated on ice to block endocytosis, and then incubated with antiserum specific for the variant surface glycoprotein (VSG), which accounts for the overwhelming majority of the surface glycoprotein coat. A sample of cells were fixed at this timepoint (t=0) to confirm surface binding of the antibodies. The remaining cells were washed (at low temperature), after which a second sample was fixed (t=1). The remaining cells were shifted to 37 °C to reactivate endocytosis and incubated for 2 minutes to allow the antibodies to enter the flagellar pocket and then be internalised by endocytosis.

**Figure 9.**
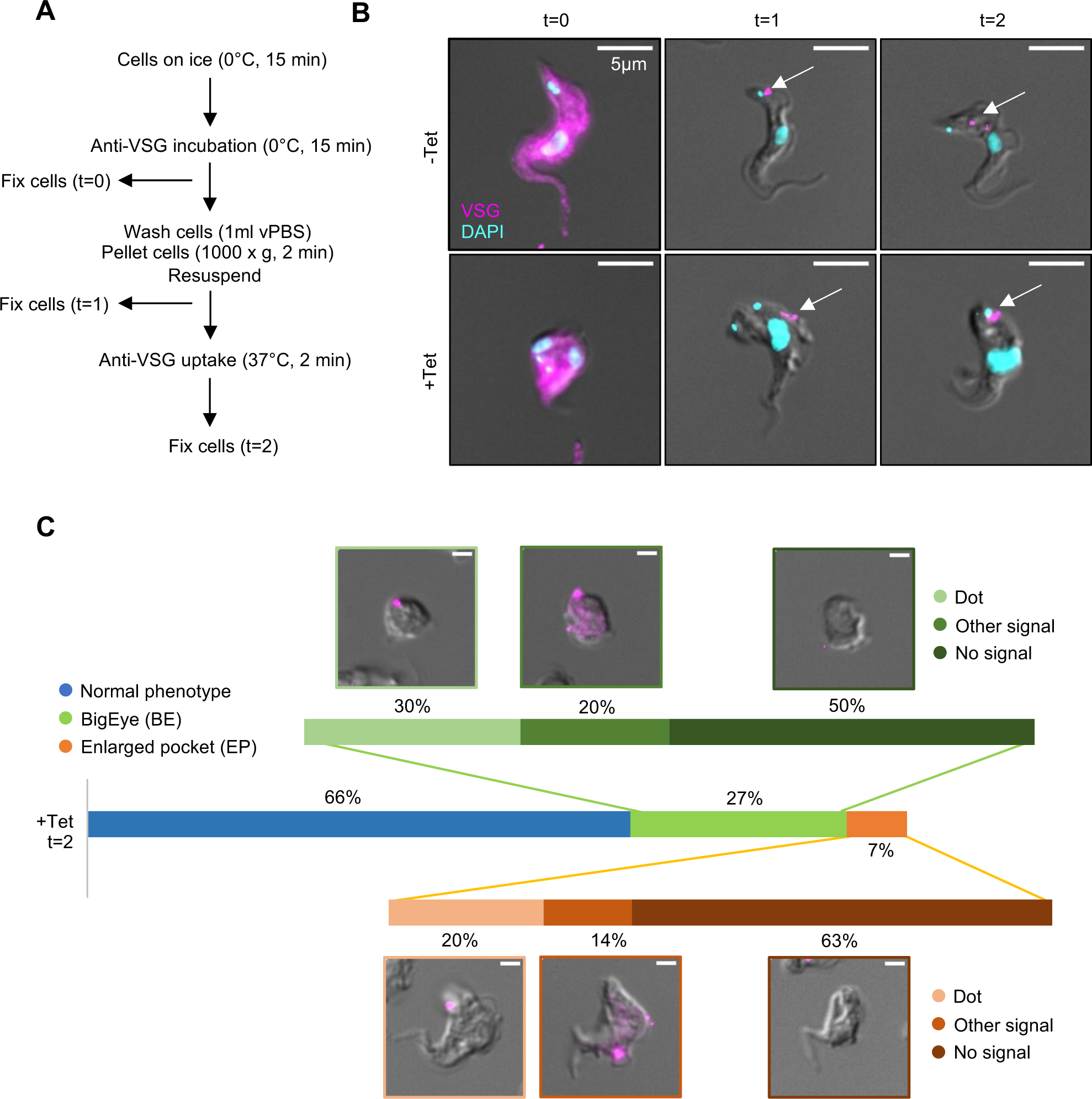
Anti-VSG does not enter the flagellar pocket of TbSmee1-depleted cells. (A) Summary of anti-VSG uptake protocol. (B) Anti-VSG is unable to enter the enlarged flagellar pocket of TbSmee1-depleted cells. In control (−Tet) cells, anti-VSG signals were observed at the cell surface (t=0), then at the location of the flagellar pocket (t=1) and finally at the endosomal/lysosomal system after allowing endocytosis (t=2) (arrows, upper panels). In TbSmee1-depleted cells (+Tet), no anti-VSG signals were observed in the enlarged flagellar pocket or in the endosomal/lysosomal system (t=1, t=2) (arrows, lower panels). Maximum intensity projections of the fluorescence channels are shown overlaid with a single DIC z-slice. (C) Quantification of anti-VSG uptake experiments. Cells from TbSmee1-depleted samples (+Tet, t=2) were manually classified into three categories (Normal phenotype, BigEye, and Enlarged pocket). The anti-VSG signal could not be observed in enlarged flagellar pockets in BigEye cells or cells with slightly enlarged flagellar pockets. Signals in these cells were classified into three categories (Dot, Other signal or No signal). Results are shown as a stacked bar chart; exemplary cells for each category are shown. Percentages represent total cell counts (408 cells) that were obtained from multiple independent experiments (n>3); each experiment included three separate clones.

The distribution of anti-VSG antibodies in control and TbSmee1-depleted cells was analysed using microscopy (Fig. 9B). In control (−Tet) cells, anti-VSG signals were observed at the cell surface (t=0), then at the location of the flagellar pocket (t=1) and finally at the endosomal/lysosomal system after allowing endocytosis (t=2) (arrows, upper panels). In TbSmee1-depleted cells (+Tet), no anti-VSG signals were observed in the enlarged flagellar pockets (t=1, t=2) (arrows, lower panels). It should be noted that there was considerable loss of anti-VSG signal between the t=0 and t=2 timepoints, and half of the TbSmee1-depleted cells displaying a phenotype had no observable signal at all. This obviously makes the conclusions from these experiments somewhat tentative.

To quantify the results, the incidence of morphologically abnormal cells was counted, and then the various distributions of anti-VSG antibodies were classified into three categories (Fig. 9C). Of the 35 % of cells in the TbSmee1-depleted population that displayed a phenotype after 24 h of RNAi, 26 % were classified as being completely rounded up (“BigEye”) while 7 % had the less-developed “enlarged pocket” state. The anti-VSG antibody distributions were grouped into three categories, respectively either a dot signal, another signal (usually a dot with other weaker labelling), and no signal. Although most of the morphologically-abnormal cells had no anti-VSG signal at all, TbSmee1-depleted cells with clearly internalised anti-VSG of the kind observed in control cells were never seen.

In summary, the inhibition of endocytosis caused by depletion of TbSmee1 prevents the internalisation of surface-bound anti-VSG antibodies, and appears to also prevent their entry into the enlarged flagellar pocket, consistent with the results obtained using ConA and BSA.

## Discussion

In this study, the hook complex component TbSmee1 was characterised in bloodstream form *T. brucei*. It is only the third hook complex protein to be characterised in this life cycle stage after TbMORN1 and Bhalin, and like them is essential for the viability of the cells in vitro (Broster Reix et al., 2021; Morriswood and Schmidt, 2015). The localisation of TbSmee1 was found to be identical to that previously documented in the procyclic life cycle stage of *T. brucei* (Figs 2, 3) (Perry et al., 2018). It is constitutively present at the shank part of the hook complex and probably also the FAZ tip, and additionally localises to the tip of the growing new FAZ in replicating cells. At the time of the preprint of this work being published, TbSmee1 was the first protein besides the DOT1 antigen and FLAM3 to be shown to localise to the groove structure associated with the tip of the growing new FAZ (Sunter et al., 2015a). A number of additional proteins present at the groove have since been identified (Smithson et al., 2022).

The analysis of truncation constructs suggested that different parts of the TbSmee1 primary structure are responsible for targeting to the two locations (Figs 4, S5). Targeting to the hook complex is mediated by the second and third predicted domains of TbSmee1. It is currently not clear whether the linker region between these two domains (aa 401-630) is also required, although the AlphaFold prediction for this region shows only unstructured loops. Nonetheless, this would be an obvious hypothesis to test in future work. Targeting to the FAZ tip appears to be mediated by the linker region (aa 161-265) between the first and second predicted domains. This is somewhat surprising, as there are few highly-conserved residues in this region (Fig. S1), and AlphaFold again predicts an unstructured loop for this region. It should be noted that while aa 161-265 have been shown to be necessary for FAZ tip targeting, they have not yet been shown to be sufficient - this again would be an obvious future experiment should this line of enquiry be pursued further. The very low observed expression levels for TbSmee1(2-160) and TbSmee1(265-640) also mean that the negative results obtained with these constructs should be treated with some caution. Given that it is unknown whether Smee1 can self-interact, the presence of the endogenous protein in these targeting experiments is an additional factor that could be addressed in follow-up work. It is unclear what role phosphorylation of TbSmee1 plays in its localisation, but given that it is both a substrate and putative direct binding partner of the mitotic kinase TbPLK, it is possible that its phosphorylation status may be related to its localisation to the groove structure.

Phenotypic characterisation of TbSmee1 showed that its depletion results in the enlargement of the flagellar pocket with concomitant effects on the entry of macromolecular cargo (Figs 5-7). The effects on the entry of macromolecular cargo are identical to those seen following TbMORN1 and Bhalin depletion (Broster Reix et al., 2021; Morriswood and Schmidt, 2015). A clear limitation of all the TbSmee1 RNAi experiments was the inability to specifically isolate cells with reduced levels of TbSmee1. As the cells were being assayed very soon after the onset of the growth defect, this meant that all assays were being conducted against a background of cells exhibiting different and often much lower levels of TbSmee1 depletion. This could be addressed in future experiments either by using a fluorescent tag on TbSmee1 to enable sorting of the subpopulation of depleted cells, or by synchronising the cells prior to the induction of RNAi. Nonetheless, there was still extremely good agreement between the immunoblotting, growth curve, and cell cycle analysis data.

At 18 h post-induction of RNAi, TbSmee1 protein levels were reaching their minimum but there was as yet no observable effect on cell cycle progression or population growth. From around 20-21 h post-induction, when TbSmee1 levels reached 10-15% of controls, cell cycle progression slowed, resulting in an increase in 2K1N and 2K2N cells. This was mirrored by a slowing of population growth. Systematic analysis of a number of marker proteins for the hook complex, the centrin arm, and the flagellar pocket collar showed that none were strongly affected in terms of their protein levels or localisation by the depletion of TbSmee1. This argues that the effects seen upon TbSmee1 depletion at these early timepoints are probably due to a loss of function, rather than due to structural perturbations in the hook complex.

In the previous characterisation of TbSmee1 in the procyclic form life cycle stage, depletion of TbSmee1 was found to cause structural changes to the hook complex and altered TbMORN1 distribution patterns (Perry et al., 2018). These analyses were however done after 6 days of RNAi induction, when TbSmee1 levels were already undetectable from 1 day onwards (a growth defect was observable from 3 days of RNAi onwards). It is very likely that similar changes would be observed in the bloodstream form stage at later RNAi timepoints, although the essential nature of TbSmee1 in bloodstream form cells might make this analysis difficult. Regardless, the focus in this study was the immediate and early effects of TbSmee1 depletion, which appear to be focused on the flagellar pocket.

Flow cytometry analysis indicated that the flagellar pocket enlargement began around 18 h post-induction, prior to the effects on population growth and cell cycle progression. This therefore suggests that flagellar pocket enlargement is an early and likely direct consequence of TbSmee1 depletion, with the effects on cell cycle progression coming afterwards. One plausible hypothesis to account for these observations is that the spatial problems produced by flagellar pocket enlargement impair cell cycle progression. The relative lack of cell cycle checkpoints means that “monster” cells with re-duplicated organelles are then produced. The internal pressure caused by continuing enlargement of the flagellar pocket forces the cells into an increasingly spherical shape and eventually cell viability is lost. In this context, the lack of a strong phenotype following TbStarkey1 depletion, despite evidence from electron microscopy that there is some enlargement of the flagellar pocket (Fig. S6), is puzzling. Cells with enlarged flagellar pockets in TbStarkey1-depleted populations were however not immediately obvious at the light microscopy level, so the prevalence of this phenotype might have been overestimated during image acquisition on the electron microscope.

The enlargement of the flagellar pocket was accompanied by a cargo access defect previously documented for TbMORN1 and Bhalin - small fluid-phase cargo such as 10 kDa dextran fills the enlarged flagellar pocket, while larger cargo such a ConA and BSA is unable to (Broster Reix et al., 2021; Morriswood and Schmidt, 2015). In TbSmee1-depleted cells, BSA was observed in one or more small foci next to the flagellar pocket. This was surprising, as BSA is known to traffic in the fluid phase and should not therefore be binding to the cell surface. BSA was however also observed to associate with the flagellar pocket membrane in an electron microscopy-based study of cargo uptake (Gadelha et al., 2009). In a classic radiochemical study of cargo uptake in *T. brucei,* it was observed that “at low concentration, a small adsorbtive component may become prevailing” in assays using BSA (Coppens et al., 1987). As the experiments conducted here and in the 2009 paper were using BSA well below physiological concentrations, it seems likely that these foci represent the small adsorbtive component that was indirectly observed in the biochemical assays. Therefore, in these assays, the ConA and BSA probes both report on the behaviour of surface-bound cargo, while the dextran reports on fluid-phase uptake.

As ConA is not a physiological cargo, the entry of anti-VSG antibodies into the flagellar pocket was also investigated. These assays turned out to be unexpectedly difficult, owing to much of the signal being lost during the wash steps. This may be related to the recently-reported high rate of VSG shedding (Garrison et al., 2021). Thus, the conclusions obtained here are necessarily tentative. Nonetheless, the t=0 and t=2 timepoints would appear to match expectations: in control cells, the anti-VSG antibodies are taken up by endocytosis, while in TbSmee1-depleted cells any remaining signal was found outside the area of the enlarged flagellar pocket and apparently still on the cell surface. It would be informative to repeat these experiments using a defined amount of affinity-purified antibody in order to minimise the need for washing, but this is outside the scope of this study.

When the TbMORN1 phenotype was characterised, this was the first time that a protein had been shown to play a role in the entry of surface-bound cargo into the flagellar pocket. Subsequent characterisation of Bhalin and the work here on TbSmee1 shows that this cargo entry defect appears to be a consistent effect after enlargement of the flagellar pocket. Furthermore, the fact that the same cargo entry defect is seen upon depletion of clathrin suggests that this is a previously-unobserved feature of all flagellar pocket enlargement (“BigEye”) phenotypes. Therefore, active endocytosis appears to be required for the entry of surface-bound cargo into the flagellar pocket.

It should be noted that the results obtained in the clathrin RNAi experiments contradict previously-published observations, where ConA was claimed to accumulate inside the enlarged flagellar pocket of clathrin-depleted cells (Allen et al., 2003). Re-examination of previously-published data from clathrin RNAi cells suggests however that the effect has always been present, with ConA signal (and indeed other cargoes) absent from the flagellar pocket interior and predominantly concentrated in one or more foci just outside it (see for example Fig. S4 in (Allen et al., 2003), and Fig. S2 in (Zoltner et al., 2015)). This oversight is understandable, given that the focus in these clathrin papers was on whether or not cargo endocytosis was occurring, and not whether the cargo was able to access the flagellar pocket (Mark Field, personal communication). Importantly, it also means that similar observations have been made by (at least) two independent groups.

It has been known for some time that surface-bound cargo in the lumen of the flagellar pocket is associated with the membrane underlying the microtubule quartet (Brickman and Balber, 1990; Gadelha et al., 2009). It has also previously been shown that upon inhibition of endocytosis, tomato lectin, wheat germ agglutinin, and BSA accumulate on this part of the flagellar pocket membrane (Gadelha et al., 2009). What is intriguing about the observations with TbSmee1 and clathrin RNAi cells here is that they show that fluid phase cargo (dextran) is still able to enter the flagellar pocket even when endocytosis is inhibited, so the effects are specific to surface-bound cargo. Given that the ConA and BSA material becomes concentrated into one or more foci that do not overlap with the dextran signal (Figs 7,8), this suggests that the majority of cargo is not able to enter the flagellar pocket lumen at all when endocytosis is inhibited.

This begs two important questions: why is it that an inhibition of endocytosis apparently hinders the entry of surface-bound cargo into the flagellar pocket? And why does the depletion of hook complex proteins - which are spatially removed from the actual sites of clathrin-coated vesicle formation - cause an endocytosis defect? The prevailing model for several years has been that hydrodynamic flow, driven by flagellar motility, is responsible for sorting surface-bound cargo to the posterior end of the cell where the flagellar pocket is located (Engstler et al., 2007). Once there, the surface-bound material enters the flagellar pocket through a narrow channel where the flagellar membrane and the flagellar pocket neck membrane are not as closely apposed (Gadelha et al., 2009). The results here suggest that the posterior sorting mechanism still functions when endocytosis is inhibited, but that the transit of surface-bound material through the channel and into the flagellar pocket might be impeded.

In this context, it is worth considering the somewhat confusing anti-VSG uptake results obtained at the t=1 timepoint in control cells, where the anti-VSG signal came from a single spot adjacent to the kinetoplast. As a temperature block was in place to prevent endocytosis, this would at first sight suggest that the anti-VSG antibodies can enter the flagellar pocket in the absence of endocytosis, and that the entire hypothesis proposed here is invalid. Given however that the flagellar pocket is less than 1 µm across, it is possible that there is insufficient resolution to distinguish the entrance to the flagellar pocket from the flagellar pocket itself in control/wild-type cells. In this interpretation, it is only when the flagellar pocket is expanded due to the BigEye phenotype that these two localisations can be resolved at normal widefield resolution. Higher-resolution imaging of these cells is therefore an important target for follow-up work.

An earlier hypothesis for hook complex function was that it was somehow maintaining the integrity of the channel in the flagellar pocket neck. This remains possible, and determining the exact site of cargo blockade, as well as imaging the microtubule quartet, will be key goals for future work. Nevertheless, the fact that the same phenotype is obtained following depletion of clathrin suggests that the hook complex is instead indirectly affecting endocytosis. This could potentially be by affecting either the localisation or activity of a number of lipid kinases, at least two of which are known to localise to the hook complex (Dean et al., 2017; Demmel et al., 2014). The activity of these enzymes could then be licensing the flagellar pocket membrane for endocytosis, for instance by generating the essential endocytic cofactor phosphatidylinositol-(4,5)-bisphosphate. Subsequent internalisation of membrane by endocytosis would then assist to pull in more membrane currently at the flagellar pocket entrance. How this would be integrated with the activity of the exocytic pathway is, however, unclear.

Thus, while hydrodynamic flow may be responsible for concentrating cargo at the entrance to the flagellar pocket, endocytic activity seems required for the entry of the cargo into the flagellar pocket itself, and endocytosis may assist or be responsible for pulling material in through the channel. Exploring these mechanisms is likely to be a fascinating area for future enquiry.

## Materials and Methods

### Recombinant protein expression and purification

The TbSmee1(1-400) open reading frame was amplified from *Trypanosoma brucei brucei* strain Lister 427 genomic DNA by PCR. The PCR product was ligated into the p3NH expression vector, which encodes an N-terminal His6 tag, using sequence and ligation-independent cloning (Li and Elledge, 2012). The plasmid was used to transform *E. coli* strain Rosetta II (DE3)pLysS by heat shock, and individual colonies were subsequently grown at 37 °C in the presence of 100 μg/ml kanamycin to an OD600 ∼ 0.8–1.0. Recombinant protein expression was induced by the addition of 50 μM IPTG, and the cells were then incubated overnight at 20 °C with shaking. Cells were harvested by centrifugation (5000× g for 30 min). The pooled pellet from 6 L of cell culture was resuspended in 300 ml of lysis buffer (50 mM Hepes pH 7.0, 500 mM NaCl, 20 mM imidazole, 5% glycerol, 0.5% Triton-X, 1 mM TCEP, 200 mM PMSF and protease inhibitor cocktail). Pellet emulsions were first homogenised by mixing on ice using a T 10 basic Ultra-Turrax dispersing instrument (IKA). Final lysis was then achieved with sonication on ice, using 3 cycles of 3 min at 50% strength. Lysates were clarified by centrifugation (18,000× g, 45 min, 4 °C). The lysates were added to a HiTrap Chelating HP 5 ml column (GE Healthcare) equilibrated with buffer A (20 mM Hepes pH 7, 300 mM NaCl, 40 mM imidazole, 2% glycerol, 1 mM TCEP), and eluted with a 100% step gradient of buffer B1 (20 mM Hepes pH 7, 300 mM NaCl, 400 mM imidazole, 2% glycerol, 1 mM TCEP). Selected peak fractions were examined by SDS-PAGE for protein content and purity. Fractions containing a dominant band at approximately 46 kDa were pooled and concentrated using Amicon Ultra centrifugal filter units with 10K pore size (MerckMillipore) according to the manufacturer’s instructions. The His6 tag was removed by 3C protease during overnight dialysis in dialysis buffer (20 mM Tris-HCl pH 7, 300 mM NaCl, 2% glycerol, 1 mM DTT). Significant losses were incurred during this step. The TbSmee1(1-400) was applied to a previously equilibrated HiTrap Chelating HP 5 ml column charged with 50 mM CoCl_2_ and coupled to a GSTrap HP 1 ml column (both GE Healthcare). Buffer A was used for equilibration. TbSmee1(1-1400) was mostly collected from the flow-through and a few initial collected fractions. Selected peak fractions were examined by 15% SDS-PAGE for protein content and purity. Fractions containing a dominant band at approximately 44 kDa were pooled and concentrated in Amicon Ultra centrifugal filter units (10K pore size) according to the manufacturer’s instructions. Finally, TbSmee1(1-400) concentrates were applied to a previously equilibrated HiLoad 16/600 Superdex 200 pg column (GE Healthcare) pre-equilibrated in dialysis buffer. Flow speed was adjusted to 1 ml/min and fractions of 1.5 ml were collected. Fractions corresponding to the targeted chromatographic peak were examined for protein content by 15% SDS-PAGE, pooled accordingly to their purity, concentrated, and stored at −80 °C.

### Antibody generation and affinity purification

Purified recombinant TbSmee1(1-400) was used for the generation of two polyclonal rabbit antisera (Eurogentec). Antisera (303, 304) were initially affinity-purified against the TbSmee1(1-400) antigen, but the neat antisera were later found to show high specificity and were also occasionally used. A third polyclonal antibody (508) was generated against two TbSmee1 peptides (Eurogentec) and affinity purified using the peptide antigens immobilised on a Sulfolink affinity column (ThermoFisher). Results shown were predominantly obtained using affinity-purified “303” and “304” anti-TbSmee1 antibodies; most immunoblotting data were generated using the “304” affinity-purified antibodies, as these had the lowest background in this application, while labelling was identical for “303” and “304” in immunofluorescence experiments. The results obtained with all three antibodies were consistent. The anti-Starkey1 anti-peptide rabbit polyclonal antibodies were generated and affinity purified in the same way.

### Antibodies

The following primary antibodies have been described previously: rabbit anti-TbMORN1 (Morriswood et al., 2013), rabbit anti-TbBILBO1 (Esson et al., 2012), mouse anti-Ty1 (“BB2”) (Bastin et al., 1996), mouse anti-TbLRRP1 (Zhou et al., 2010), mouse anti-TbCentrin4 (“6C5” (Ikeda and de Graffenried, 2012)), mouse anti-TbCentrin2 (“2B2H1”) (de Graffenried et al., 2013), mouse anti-TbFAZ1 (“L3B2”) (Kohl et al., 1999), mouse anti-PFR1,2 (“L13D6”) (Kohl et al., 1999), mouse anti-FAZ filament (“DOT1”)(Woods et al., 1989), rabbit anti-VSG(221) (Batram et al., 2014). The following antibodies came from commercial sources: goat anti-rabbit(IRDye800CW) (LI-COR), goat anti-mouse(IRDye680LT) (LI-COR), goat anti-rabbit and anti-mouse antibodies conjugated to AlexaFluor dyes (Molecular Probes).

### Cell culture

Wildtype Lister 427 (monomorphic) BSF cells were cultured in HMI-9 medium (Hirumi and Hirumi, 1989) supplemented with 10 % foetal bovine serum (FBS), 100 U/ml penicillin and 0.1 mg/ml streptomycin at 37°C and 5% CO_2_. The SM (single marker) cells (Wirtz et al., 1999) were cultured in the presence of G418 (2.5 μg/ml). Population density was monitored using a Z2 Coulter Counter (Beckman Coulter), and kept below 2×10^6^ cells/ml.

### Generation of transgenic cell lines

The Ty1-TbSmee1 endogenous replacement cell line was generated by transfection of 427 cells with a targeting fragment containing 285 bp of the TbSmee1 5’ untranslated region (UTR), a blasticidin resistance gene, the intergenic region of alpha- and beta-tubulin, the sequence for a triple Ty1-tag and the first 399 bp of the TbSmee1 open reading frame (ORF) without the start codon. Clones were selected by growth in medium containing 5 μg/ml blasticidin. RNAi target sequences were chosen using RNAit (Redmond et al., 2003). TbSmee1 RNAi cells were generated by cloning the RNAi target sequence into the pGL2084 plasmid (Jones et al., 2014) and then transfecting 2T1 cells with the linearised plasmid (Alsford et al., 2005). Clones were selected by growth in medium containing phleomycin (2.5 μg/ml) and hygromycin (5 μg/ml). TbStarkey1 RNAi cells and TbCHC RNAi cells were generated by cloning the relevant RNAi target sequence into the p2T7_TAblue plasmid (Alibu et al., 2005) and then transfecting SM cells with the linearised plasmid. Clones were selected by growth in medium containing G418 (2.5 μg/ml) and hygromycin (5 μg/ml). Ty1-TbSmee1 truncation constructs were cloned into the pLew100_v5-Hyg plasmid using in vivo assembly (Watson and Garcia-Nafria, 2019). SM cells were transfected with the linearised plasmid. Clones were selected by growth in medium containing G418 (2.5 μg/ml) and hygromycin (5 μg/ml). For transfection >2.5 × 10^7^ cells of the parental strain were washed and resuspended in 100 μl transfection buffer (90 mM Na_2_PO_4_, 5 mM KCl, 0.15 mM CaCl_2_, 50 mM HEPES [4-(2-hydroxyethyl)-1-piperazineethanesulfonic acid]-NaOH, pH 7.3) containing 10 μg DNA and transfected by electroporation using an AMAXA Nucleofector® Device (Lonza) with program “X-001 free choice”. Transfected cells were incubated in 50 ml HMI-9 medium without selection overnight. The next day, drug selection was applied and clones were selected by limiting dilution. At least three separate clones of all cell lines were isolated to control for biological variability. Integration of targeting fragments at endogenous loci or the presence of Ty1-TbSmee1 truncation constructs in the genome were confirmed by PCR analysis of genomic DNA. Genomic DNA was isolated using a DNeasy Bloody & Tissue kit (QIAGEN) and relevant products were amplified by PCR. All cloned constructs used for cell line generation had their DNA sequence confirmed by sequencing.

### Immunoblotting

For generation of dephosphorylated whole-cell lysates, cells were harvested by centrifugation (1000xg, 10 min) and the cell pellet was resuspended in 1 ml vPBS (PBS, 46 mM sucrose, 10 mM glucose) containing EDTA-free protease inhibitors (Roche). The washed cells were pelleted by centrifugation (750xg, 4 min). The cells were then resuspended in lysis buffer (0.5% IGEPAL, 0.1M PIPES-NaOH pH 6.9, 2 mM EGTA, 1 mM MgCl2, 0.1 mM EDTA, EDTA-free protease inhibitor cocktail) to a final concentration of 4×10^5^ cells/μl and incubated for 15 min at RT on an orbital mixer to allow dephosphorylation to occur. SDS-loading buffer was then added to a final concentration of 2×10^5^ cells/μl, the samples were further denatured by boiling (100°C, 10 min), and then stored at −20°C. SDS-PAGE was carried out using a Mini-Protean Tetra Cell (Bio-Rad), and protein transfer to nitrocellulose membranes using a Mini-Trans blot cell (Bio-Rad). Protein transfer and equal loading was confirmed using REVERT total protein stain (LI-COR) according to the manufacturer’s instructions. Membranes were blocked using blocking buffer (PBS, 0.3% Tween 20, 10% milk)(30 min, RT, rocker). The membranes were then incubated in primary antibodies diluted in blocking buffer (1 h, RT, roller). After three washes in immunoblot buffer (PBS, 0.3% Tween 20) the membranes were incubated with IRDye-conjugated secondary antibodies diluted in immunoblot buffer (1 h, RT, roller). After another three washes in immunoblot buffer, the membranes were visualised using an Odyssey CLx (LI-COR). Background subtraction, qualitative analysis, normalisation relative to total protein stain, and quantification were carried out using Image Studio Lite 5.2 and Empiria Studio 1.1 (LI-COR). Dot plots were generated using the web app PlotsOfData (Postma and Goedhart, 2019).

### In vitro phosphatase assays

427 BSF cells were grown to approximately 1.5×10^6^ cells/ml and harvested by centrifugation (1,000xg, 10 min, 4 °C). The cells were washed in 1 ml ice-cold wash buffer (0.1M PIPES-NaOH pH 6.9, 2 mM EGTA, 1 mM MgCl2, 0.1 mM EDTA, phosphatase Inhibitor Cocktail 2) and pelleted by centrifugation (750xg, 3 min, 4 °C). For extraction of the cytoskeletons, the cell pellet was resuspended in 1 ml ice-cold extraction buffer (0.5% IGEPAL, 0.1M PIPES-NaOH pH 6.9, 2 mM EGTA, 1 mM MgCl2, 0.1 mM EDTA, EDTA-free protease inhibitor cocktail, phosphatase Inhibitor Cocktail 2) and incubated for 15 min on ice. The cell suspension was inverted every 5 min. The cytoskeletons were separated from the cytoplasm by centrifugation (750xg, 3 min, 4 °C). The cytoskeletons were washed with 0.5 ml ice-cold 1x NEBuffer for PMP supplemented with 1 mM MnCl_2_ and pelleted by centrifugation (750xg, 3 min, 4 °C). After resuspension in 375 μl ice-cold 1x NEBuffer for PMP supplemented with 1 mM MnCl_2_, all following samples were taken from this stock. An input sample (0 min) of 40 μl was taken and added to 20 μl of SDS-loading buffer. Two control samples were taken, 0.4 μl Phosphatase Inhibitor Cocktail 2 was added, and incubated on ice and at 26°C, respectively, for 20 min. 20 μl of SDS-loading buffer was then added. To assay for dephosphorylation, 130 μl were taken from the stock and 1 μl lambda phosphatase (400 U) was added. This sample and the remainder from the stock were incubated at 26 °C. After 1/2/3/5/10/20 min 20 μl samples were taken from each and were added to 10 μl SDS-loading buffer. All the samples were boiled at 104 °C for 10 min and stored at −20 °C. Samples were analysed by immunoblotting.

### Fractionation

50 ml 427 BSF cells were grown to approximately 1.5×10^6^ cells/ml and then harvested by centrifugation (1000xg, 10 minutes, 4°C). The cell pellet was resuspended in 1 ml vPBS, transferred to a microfuge tube, and the cells again pelleted by centrifugation (750xg, 4 min, 4°C). The supernatant was discarded and the centrifugation step was repeated to remove remaining supernatant. The cell pellet was resuspended in 200 μl extraction buffer (0.5% IGEPAL, 0.1M PIPES-NaOH pH 6.9, 2 mM EGTA, 1 mM MgCl2, 0.1 mM EDTA, EDTA-free protease inhibitor cocktail) and incubated for 15 minutes at RT in an orbital mixer. A 5% input sample was taken (10 μl), put into a new microfuge tube and left on ice. The extracted cells were fractionated by centrifugation (3400xg, 2 minutes, 4°C) and the supernatant transferred to a fresh microfuge tube and the exact volume noted. The tube containing the extracted cells was centrifuged again at the same settings and this second residual supernatant discarded. The cytoskeleton pellet was then resuspended in 200 μl extraction buffer. 5% samples of supernatant and pellet fractions were taken and analysed by immunoblotting. Equal fractions of I, SN, P were loaded in each lane (I ∼ 1.4×10^6^ cells).

### Preparation of samples for immunofluorescence microscopy

Coverslips were washed in 70% ethanol and then incubated with 0.01% poly-L-lysine in a 24-well plate (>20 min, RT) and left to dry. 2×10^6^ cells were taken per coverslip and transferred to 15 ml Falcon tubes. The cells were pelleted by centrifugation (1000xg, 1 min per ml of liquid, RT) in a swing-bucket centrifuge. The supernatant was removed, and the cell pellet was gently resuspended in 1 ml ice-cold vPBS + Complete. The cells were again pelleted by centrifugation (1000xg, 2 min, RT) and subsequently resuspended in 1 ml ice-cold vPBS + Complete and directly added to the coverslips. The cells were attached to the coverslips by centrifugation (1000xg, 1 min, RT), and attachment was confirmed visually. The attached cells were then incubated in 1 ml ice-cold extraction buffer (0.5% IGEPAL 0.1M PIPES-NaOH pH 6.9, 2 mM EGTA, 1 mM MgCl2, 0.1 mM EDTA, EDTA-free protease inhibitor cocktail) (5 min, on ice). The extracted cells were washed two times with 1 ml vPBS + Complete and then fixed in 1 ml ice-cold 99.9% methanol (30 min, −20°C). The fixed cells were rehydrated using 1 ml PBS. The coverslips were blocked in 1 ml 3% BSA in PBS (30 min, RT), and sequentially incubated with clarified primary and secondary antibodies diluted in PBS (1 h, RT, humidified chamber for each) with three PBS washing steps (3 x 5 min, RT, rocker) after each incubation. After the final wash, glass slides were cleaned with 70% ethanol and a spot of DAPI-Fluoromount G (Southern Biotech) was placed on the surface. The coverslips were rinsed in ddH_2_O, carefully dried, and then mounted. For preparation of samples for SIM imaging, different coverslips (high precision, No. 1.5 H) were used. Cells were harvested as above, washed, and fixed in 4% paraformaldehyde solution (10 min, ice then 30 min, RT). The fixed cells were washed, attached to coverslips, permeabilised (0.25% Triton X-100 in PBS; 5 min, RT), and then labelled and mounted as described above.

### Fluorescence microscopy

Images were acquired using a DMI6000B widefield microscope (Leica Microsystems, Germany) with a HCX PL APO CS objective (100x, NA = 1.4, Leica Microsystems, Germany) and Type F Immersion Oil (refractive index = 1.518, Leica Microsystems, Germany). The microscope was controlled using LAS-X software (Leica). Samples were illuminated with an EL6000 light source (Leica) containing a mercury short-arc reflector lamp (HXP-R120W/45C VIS, OSRAM, Germany). Excitation light was selected by using Y3 (545/25 nm), GFP (470/40 nm), and A4 (360/40 nm) bandpass filter cubes (Leica Microsystems, Germany). The power density, measured at the objective focal plane with a thermal sensor head (S175C, Thorlabs), was respectively 0.749 ± 0.086, 0.557 ± 0.069, 0.278 ± 0.076 W/cm^2^ for the three filters. Emitted light was collected at ranges of 605/70 (Y3), 525/50 nm (GFP), and 470/40 nm (DAPI) respectively. The individual exposure times and camera gains were adjusted according to the different samples. RNAi samples (control and depleted) were imaged using identical settings. Differential interference contrast (DIC) was used to visualise cell morphology. 3D recording of each field of view was obtained using 40 Z-slices (step size = 0.21 µm). Fields of view were selected in the DIC channel in order to blind the user to the fluorescence signal and subjectively select for cells with optimum morphology. Images were captured using a DFC365 FX monochrome CCD camera (Leica, 6.45 µm pixel size). SIM images were acquired using an Elyra S.1 SIM microscope (Zeiss) and ZEN software (Zeiss).

### Fluorescence microscopy image processing and analysis

Processing was carried out using FIJI (Schindelin et al., 2012) and a custom macro for the generation of maximum-intensity z-projections with single DIC z-slices overlaid. Overlaps between two proteins were confirmed in individual z-slices (thickness: 210 nm). The plugin ScientiFig was utilised to create the collage and adding the inserts (Aigouy and Mirouse, 2013). For correlation analysis between TbSmee1 and other flagellar pocket collar and/or hook complex associated proteins, 1K1N cells were selected using both DIC and DAPI channels. 2D sum slices projections were prepared for each stack of both green and red channels. The projections were clipped to 8-bit depth and a convoluted background subtraction was applied. All resultant individual 2D images, without channel overlay, were analysed pairwise to check for intensity-based correlation based on Zhang & Cordelières (Zhang and Cordelières, 2016). The Spearman’s rank correlation results were further computed into Microsoft Excel sheets and analysed using R version 4.1.2 (R Core Team, 2022) in the environment RStudio 2021.09.1.372 (RStudio Team, 2020). The packages used for all descriptive analysis and plot generation were ggplot2 (Wickham, 2016), openxlsx (Schauberger and Walker, 2022), and psych (Revelle, 2023). All Fiji/ImageJ macros and R scripts for correlation analysis were written by Alyssa Borges and are available upon request.

### Growth curves

RNAi cells were seeded at the required starting concentration in a volume of 22 ml and divided into two 10 ml aliquots in separate flasks. Tetracycline was added to a final concentration of 1 μg/ml in one flask to induce RNAi, and refreshed every 24 h. The population density of the control and induced cells was measured every 24 h over a time course of 72h, or every hour over a time course of 8 h, using a Z2 Coulter Counter (Beckman Coulter). Depletion of the target protein was confirmed in every experiment by immunoblotting of whole-cell lysates.

### Cell division cycle analysis

An aliquot of 10^6^ cells was taken, and the cells were fixed directly in media by addition of isothermal 25% glutaraldehyde to a final concentration of 2.5% (10 min, 37°C, gentle mixing). The cells were then pelleted by centrifugation (750xg, 10 min). The cell pellet was resuspended in 0.5 ml 2.5% glutaraldehyde in PBS, transferred to a microfuge tube, and incubated at RT (30 min, gentle mixing). The cells were pelleted again by centrifugation (750xg, 4 min), and the cell pellet resuspended in 500 μl PBS. The cells were then added to the coverslips inside the 24-well plate and attached by centrifugation (1000xg, 1 min, RT). The coverslips were mounted on poly-L-lysine-coated slides using DAPI-Fluoromount G (Southern Biotech). Imaging was as described for fluorescence microscopy above, using DAPI and DIC channels only. Cell division cycle stages (1K1N, 2K1N, 2K2N) were manually quantified from maximum intensity projections of the DAPI signal overlaid with single DIC z-slices. Depletion of the target protein at each timepoint was confirmed by immunoblotting of whole-cell lysates from the same experiments.

### Preparation and imaging of electron microscopy samples

Induced and uninduced RNAi cells were grown for 24 h to a density of 1-2×10^6^ cells/ml in 50 ml medium and harvested by centrifugation (1000xg, 10 min, RT). The supernatant was removed to 4 ml and 4 ml FBS was added. The cells were pelleted again (1000xg, 10 min, RT) and the supernatant was removed to 200 μl. The cells were resuspended in the supernatant, and the suspension was then transferred to a PCR tube and pelleted by centrifugation (1,600xg, 10s, RT). The cells were then transferred into a carrier with a closed lid to avoid air inclusions. High pressure freezing (HPF) was started immediately (Leica EM HPM100). After HPF the samples were transferred to an AFS (Leica EM AFS2) for freeze substitution and progressive lowering of temperature. Low temperature embedding and polymerisation of Epon raisin (DDSA, MNA, Epon812, 2,4,6 Tris(dimethylaminomethyl(phenol))) were then carried out. Ultra-thin cuts (60 nm) were carried out with an ultramicrotome (Leica EM UC7/FC7) and were placed on slotted grids. For contrasting they were incubated in 2 % uranyl acetate for 8 min. Afterwards the grids were washed 3x in ddH_2_O (boiled to remove CO_2_) and incubated for 5 min on 50 % Reynold’s lead citrate in a petri dish with NaOH tablets. The grids were again washed 2x in ddH_2_O. A 200 kV transmission electron microscope (Jeol, JEM-2100) with a TemCam F416 4k x 4k camera (Tietz Video and Imaging Processing Systems) and EMMenu 4.0.9.31 software (Tietz) were used. Uninduced control cells were viewed at a magnification of 12,000x, induced cells were viewed at a magnification of 8,000x.

### Measurement of flagellar pocket enlargement

Induced and uninduced RNAi cells at a concentration of ∼5×10^6^ cells/ml were harvested by centrifugation (1000xg, 4 °C). The cells were resuspended in 45 μl ice-cold vPBS + protease inhibitors and incubated on ice to block endocytosis (10 min, ice). 5 µl labelled dextran (10 kDa, 50 mg/ml stock) was added and mixed by flicking. The mixture was then incubated to allow dextran to enter and fill the flagellar pocket (15 min, on ice, dark). At the end of the incubation, 1 ml ice-cold vPBS + protease inhibitors was added and the cells pelleted by centrifugation (750 xg, 2 min, 4 °C). The cell pellet was resuspended in 0.5 ml vPBS + protease inhibitors, and the cells fixed by addition of 0.5 ml 8 % pfa and 0.1 % glutaraldehyde solution in vPBS + protease inhibitors (20 min, on ice then 60 min, RT). The fixed cells were pelleted by centrifugation (750 x*g*, 2 min) and washed twice with 1 ml vPBS + protease inhibitors. The dextran signal was immediately measured by flow cytometry using a FACSCalibur (BD Biosciences) running CellQuest ProTM Software (BD Biosciences). Later processing and analysis was carried out using FlowJo 10.8.1. Dot plots were created using Plots of Data (Postma and Goedhart, 2019). After flow cytometry, the remaining cells were attached to coverslips by centrifugation, mounted on clean glass slides using DAPI-Fluoromount G (Southern biotech), and imaged using fluorescence microscopy on the same day to confirm flagellar pocket labelling. Depletion of TbSmee1 was confirmed in every experiment by immunoblotting of whole-cell lysates.

### Cargo (ConA, BSA) uptake assays

2×10^6^ cells per sample were harvested by centrifugation (1000 xg, 10 min, 4 °C) and washed in 1 ml ice-cold vPBS. The washed cells were pelleted by centrifugation (750 xg, 2 min, 4 °C), and resuspended in 100 µl ice-cold vPBS. The cells were then incubated at low temperature (20 min, ice) to block endocytosis. During this incubation, the labelled cargoes were prepared. The 50 mg/ml dextran aliquot (10 kDa AF488 conjugate, Molecular Probes) was mixed using a sonicator bath (10 min, 37 Hz) and then vortexed. The 0.5 mg/ml ConA aliquot (TMR conjugate, Molecular Probes) was clarified by centrifugation (11,000 xg, 10 min, 4 °C). Both cargoes were kept on ice and in the dark. At the end of the incubation, 10 µl of the 50 mg/ml dextran and 2 µl of the 0.5 mg/ml ConA were added to the chilled cells, mixed by flicking, and incubated at low temperature (15 min, on ice, in the dark). T=0 samples were quenched and fixed at this point; t=30 samples had an additional incubation to allow endocytosis of cargo (30 min, 37 °C, in the dark) before quenching and fixing. To quench samples, 1 ml ice-cold vPBS was added, and the cells were pelleted by centrifugation (750 xg, 2 min, 4 °C). The pelleted cells were resuspended in 50 µl vPBS by flicking, and then fixed by addition of 0.5 ml ice-cold fix solution (4% paraformaldehyde solution, 0.1% glutaraldehyde in vPBS) (20 min, on ice, in the dark, then 30 min, RT, in the dark). The fixed cells were pelleted by centrifugation (750 xg, 2 min, RT), washed in 2 ml vPBS, resuspended in 1 ml vPBS, and then attached to poly-L-lysine-coated coverslips by centrifugation. Coverslips were mounted on glass slides using DAPI-Fluoromount G (Southern Biotech) and imaged immediately. Assays using BSA (AlexaFluor555 conjugate, Molecular Probes) were done in the same way. The 20 mg/ml BSA aliquot was clarified by centrifugation (750 xg, 1 min, RT)), and 3 µl was added to the cells/dextran mixture. Depletion of TbSmee1 was confirmed by immunoblotting of whole-cell lysates in every experiment.

### Anti-VSG uptake assay

2×10^6^ cells per sample were harvested by centrifugation (1000 xg, 1 min per ml of liquid + 1, 4 °C) and washed in 1 ml ice-cold vPBS. The washed cells were pelleted by centrifugation (1000 xg, 3 min, 4 °C), resuspended in 50 µl ice-cold vPBS and incubated at low temperature to block endocytosis (20 min, ice). During incubation, the primary and secondary antibodies were clarified by centrifugation (10,000 xg, 10 min, 4 °C). The primary antibody (anti-VSG antiserum) was diluted 1:50 in ice-cold vPBS. The secondary antibody was diluted to required concentration (1:3000) in 3 % BSA in PBS. Both were kept on ice in the dark until used. At the end of the incubation, 50 µl of diluted anti-VSG was added to the cells and quickly mixed by flicking, followed by an incubation to allow binding (15 min, ice). To fix cells for t=0, 0.5 ml ice-cold vPBS and 0.5 ml ice-cold 8 % pfa in vPBS were sequentially added at the end of the incubation and kept on ice. For timepoints t=1 & t=2, 1 ml ice-cold vPBS was added. The cells were washed by centrifugation (1000 xg, 2 min, 4 °C) and resuspended in 1 ml ice-cold vPBS. To fix t=1, 350 µl of ice-cold 16% pfa was added and the cells were kept on ice. The cells for t=2 were incubated (2 min, 37°C) to allow redistribution, flagellar pocket entry, and endocytosis of the anti-VSG. 350 µl of ice-cold 16 % pfa was then added to fix the cells. All three timepoints were incubated on ice (20 min) after addition of pfa, followed by a subsequent incubation at RT (30 min). The fixed cells were pelleted by centrifugation (1000 xg, 2 min, RT), resuspended in 1 ml RT vPBS, and attached to poly-L-lysine-coated coverslips by centrifugation (1000 xg, 1 min, RT). The attached cells were then permeabilised in 1 ml 0.25 % Tx100 in PBS (5 min, RT). The permeabilised cells were washed two times in 1 ml PBS and the coverslips with cells were placed on a drop of clarified and diluted secondary antibody (70 µl) in a humidified chamber and incubated (1h, RT, dark). The coverslips were washed 3 times in 1 ml PBS (5 min, RT, rocker, dark). After rinsing the cells in ddH_2_O, they were mounted onto a drop of DAPI-Fluoromount G (Southern Biotech) on a coverslip that was cleaned with 70% EtOH.

## Acknowledgements

Antibodies were obtained from the following sources: Cynthia He (National University of Singapore) provided the anti-Ty1 (BB2) and anti-TbLRRP1 antibodies; Chris de Graffenried (Brown University) provided the anti-TbCentrin2 (2B2H1) and anti-TbCentrin4 (6C5) antibodies; Markus Engstler provided the anti-VSG antibodies; Keith Gull (University of Oxford) provided the anti-FAZ1 (L3B2), anti-PFR1,2 (L13D6) and DOT1 antibodies. Nadine Weisert provided assistance with cloning. Philip Kollmannsberger assisted the correlation analysis. Markus Engstler provided support and advice. Balázs Szöőr (University of Edinburgh) and Mick Urbaniak (Lancaster University) provided input on the phosphatase assays. Mark Field (University of Dundee) provided extremely helpful advice and context on earlier clathrin RNAi work. Jay Bangs (University at Buffalo) provided helpful advice on the interpretation of the anti-VSG uptake data. Tim Krüger provided input and assistance with imaging. Helena Jambor (TU Dresden) advised on data visualisation. Graham Warren (Max Perutz Laboratories) hosted the preliminary experiments for this project. Susanne Kramer and Manfred Alsheimer provided feedback on the initial manuscript. Various members of the trypanosome research community provided feedback during the revision of the manuscript.

---

**Figure S1.**
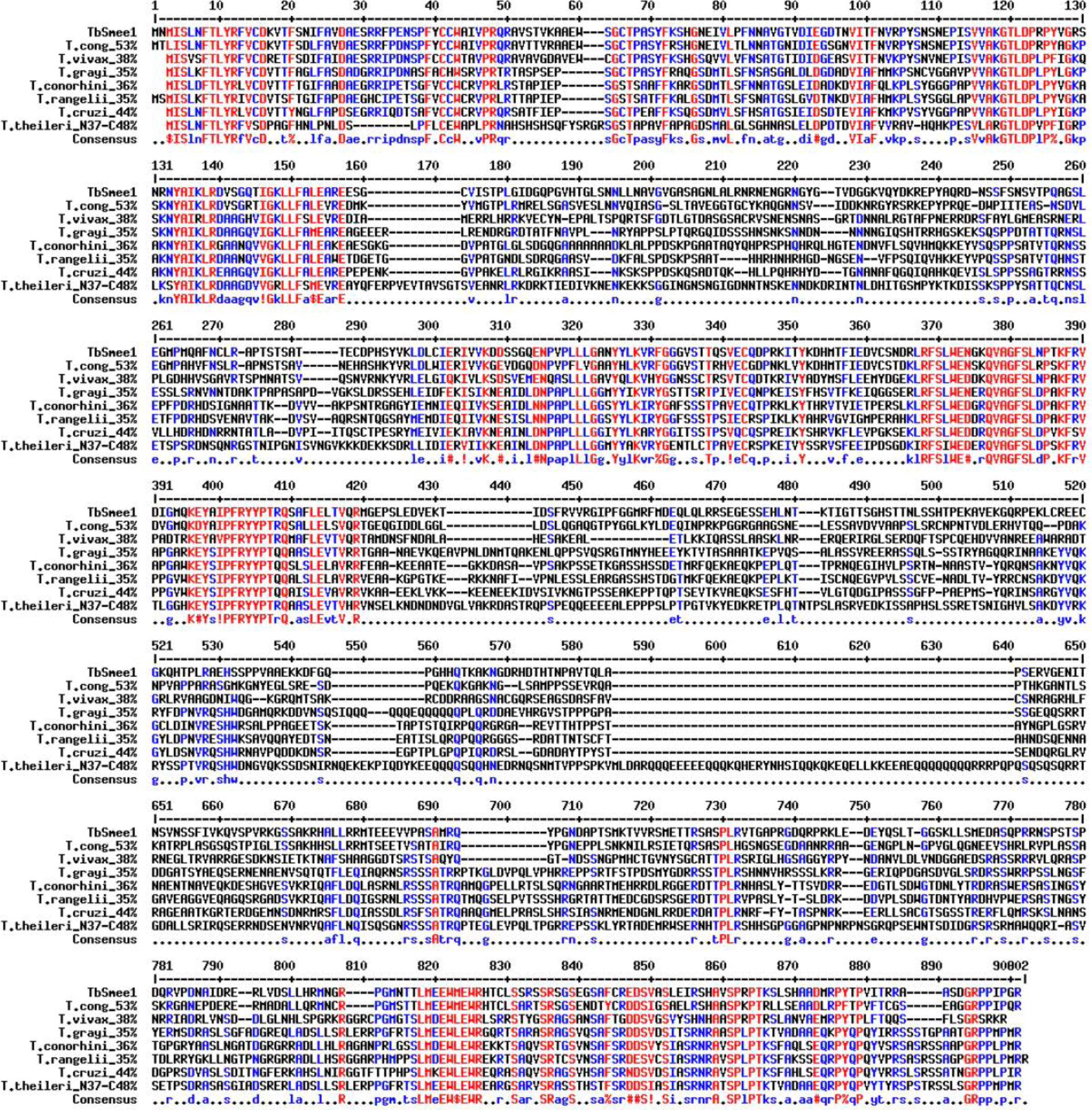
The TbSmee1 primary structure contains three blocks of highly-conserved sequence. Multiple sequence alignment of *Trypanosoma brucei* Smee1 (TbSmee1) and homologous proteins from other trypanosome species, generated using MultAlin (Corpet, 1988) with default parameters. Amino acid numbers are indicated in black numerals above the alignment. Moderately (50-90%) conserved residues are highlighted in blue. Highly-conserved (>90%) or completely conserved residues are highlighted in red. The consensus sequence is shown below the alignment. Abbreviations: TbSmee1, *Trypanosoma brucei* Smee1; T.cong, *Trypanosoma congolense*; T.vivax, *Trypanosoma vivax*; T.grayi, *Trypanosoma grayi*. T.conorhini, *Trypanosoma conorhini*; T.rangelii, *Trypanosoma rangelii*; T.cruzi, *Trypanosoma cruzi*; T.theileri, *Trypanosoma theileri*. The % sequence identity of each homologue to TbSmee1 is indicated after the name.

**Figure S2.**
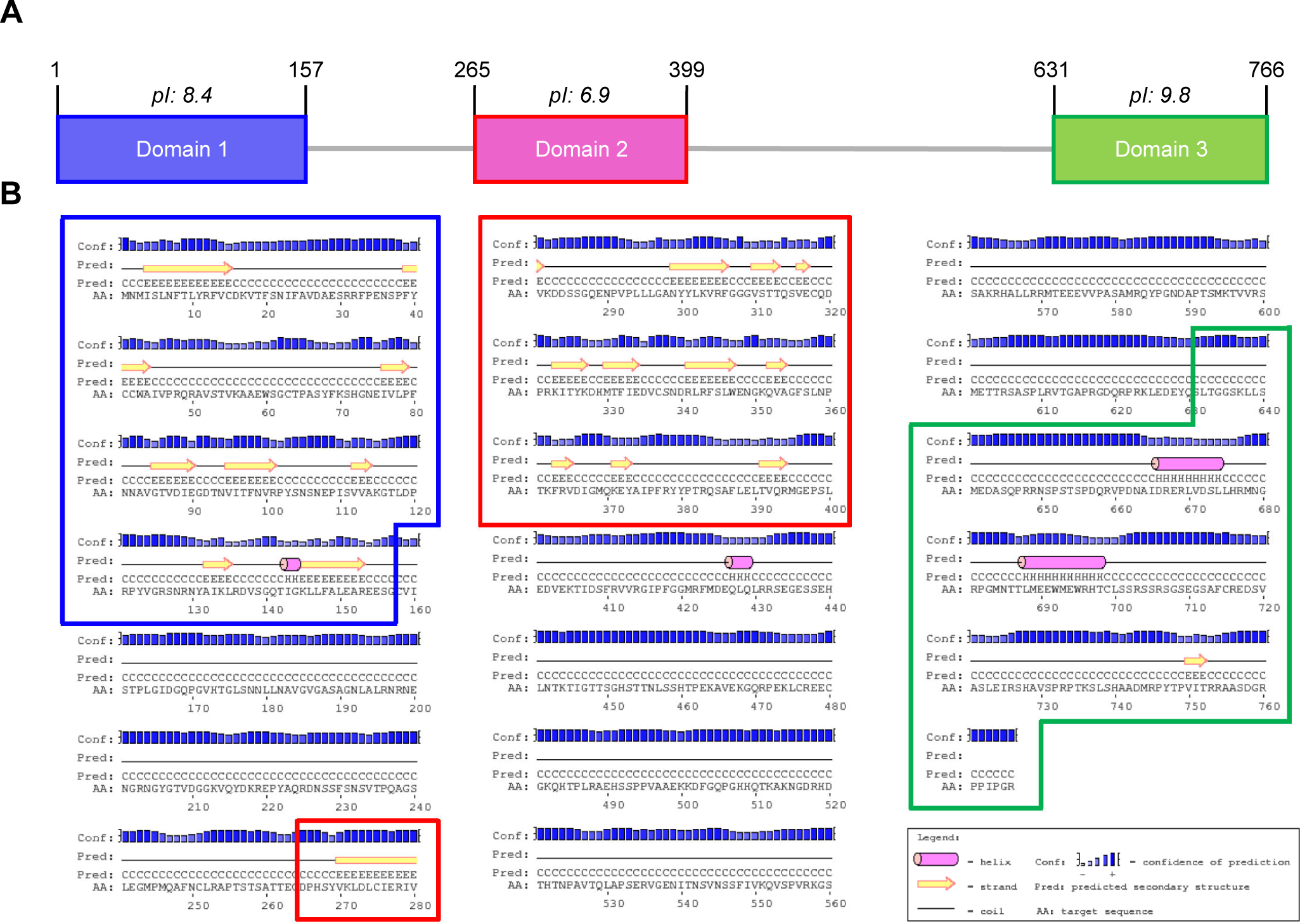
TbSmee1 contains three predicted structured domains. (A) Schematic representation of TbSmee1 with the three predicted domains (Dom1/2/3) indicated in blue, magenta, and green. The approximate amino acid coordinates in the primary structure are indicated in black above each domain. The isoelectric point (pI) of each domain is indicated above the domain. (B) TbSmee1 contains three regions of predicted secondary structure. Secondary structure prediction generated using the PSIPRED server (Buchan & Jones, 2019) with default parameters. Abbreviations and schematics are defined in the legend at the bottom right. The three predicted structured domains are outlined as blue, red, and green boxes.

**Figure S3.**
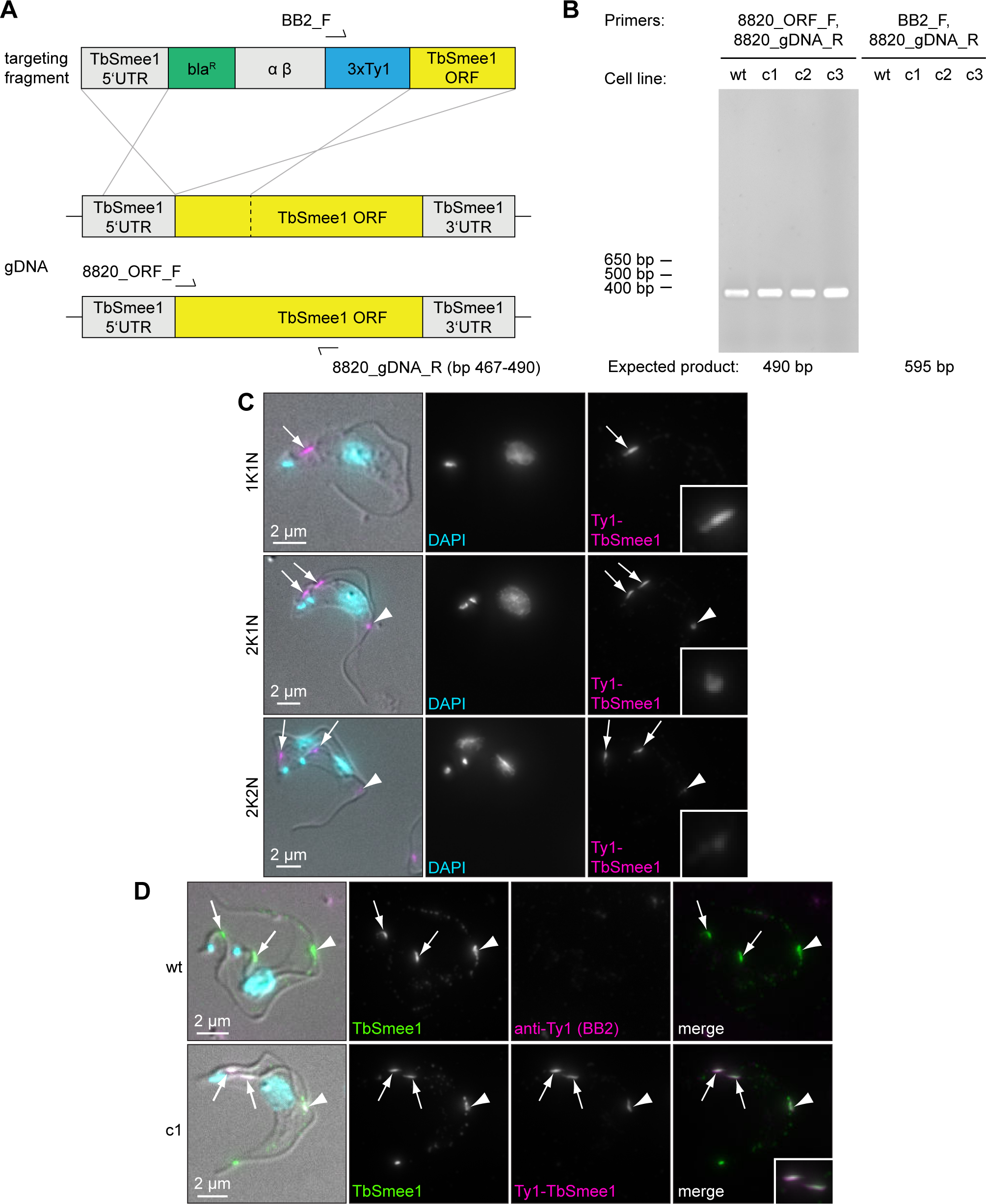
Generation of Ty1-TbSmee1 endogenous replacement cell line. (A) Schematic showing the in situ tagging procedure and annealing sites for PCR primers. The two endogenous alleles of TbSmee1 in the gDNA are shown in yellow, flanked by 5’UTR and 3’UTR sequences. *T.* brucei cells were transfected with a targeting fragment with homology arms for recombination with the 3’ end of the 5’UTR and 5’ end of the ORF. The targeting fragment contained a blasticidin resistance gene (bla^R^), the intergenic region from the alpha/beta tubulin locus (αβ) and a 3xTy1 epitope tag preceded by an ATG start codon. Homologous recombination removed the endogenous ATG start codon of the TbSmee1 ORF. (B) Confirmation of targeting fragment integration at the endogenous TbSmee1 locus by PCR analysis of genomic DNA. Genomic DNA from wild-type (wt) and candidate Ty1-TbSmee1 clones (c1, c2, c3) was analysed by PCR. Left panel: positive control using 8820_ORF_F and 8820_gDNA_R primers; 490 bp product expected. Right panel: integration test using BB2_F and 8820_gDNA_R primers. A product is only expected if the 3xTy1 sequence has integrated upstream of the TbSmee1 ORF (see primer annealing sites in panel A). Two independent experiments were carried out, each using all three separate clones. (C) Ty1-TbSmee1 displays the same localisations through the cell cycle as endogenous TbSmee1. Detergent-extracted cells were fixed with methanol and labelled with anti-Ty1 antibodies; DNA was stained using DAPI. Exemplary cells from the three main cell cycle states (1K1N, 2K1N, 2K2N) are shown. Maximum-intensity z-projections of the fluorescence channels are shown, together with a single DIC z-slice overlay. The Ty1-TbSmee1 signal is shown in magenta in the overlay and highlighted with arrows. Arrowheads indicate the Ty1-TbSmee1 present at the tip of the new FAZ. Insets show an enlarged view of the TbSmee1 signal from the hook complex or FAZ tip. Multiple (n>3) independent experiments were carried out using three separate clones. (D) The anti-Ty1 signal is specific for Ty1-TbSmee1. Wild-type (wt) and Ty1-TbSmee1 cells were extracted with detergent, fixed with methanol, and labelled with anti-TbSmee1 and anti-Ty1 antibodies. Hook complex (arrows) and FAZ tip (arrowheads) localisations are indicated. No anti-Ty1 signal was seen in wild-type cells; strong overlap between the anti-Ty1 and anti-TbSmee1 signals was seen in the Ty1-TbSmee1 cells. Maximum-intensity z-projections of the fluorescence channels are shown, together with a single DIC z-slice overlay.

**Figure S4.**
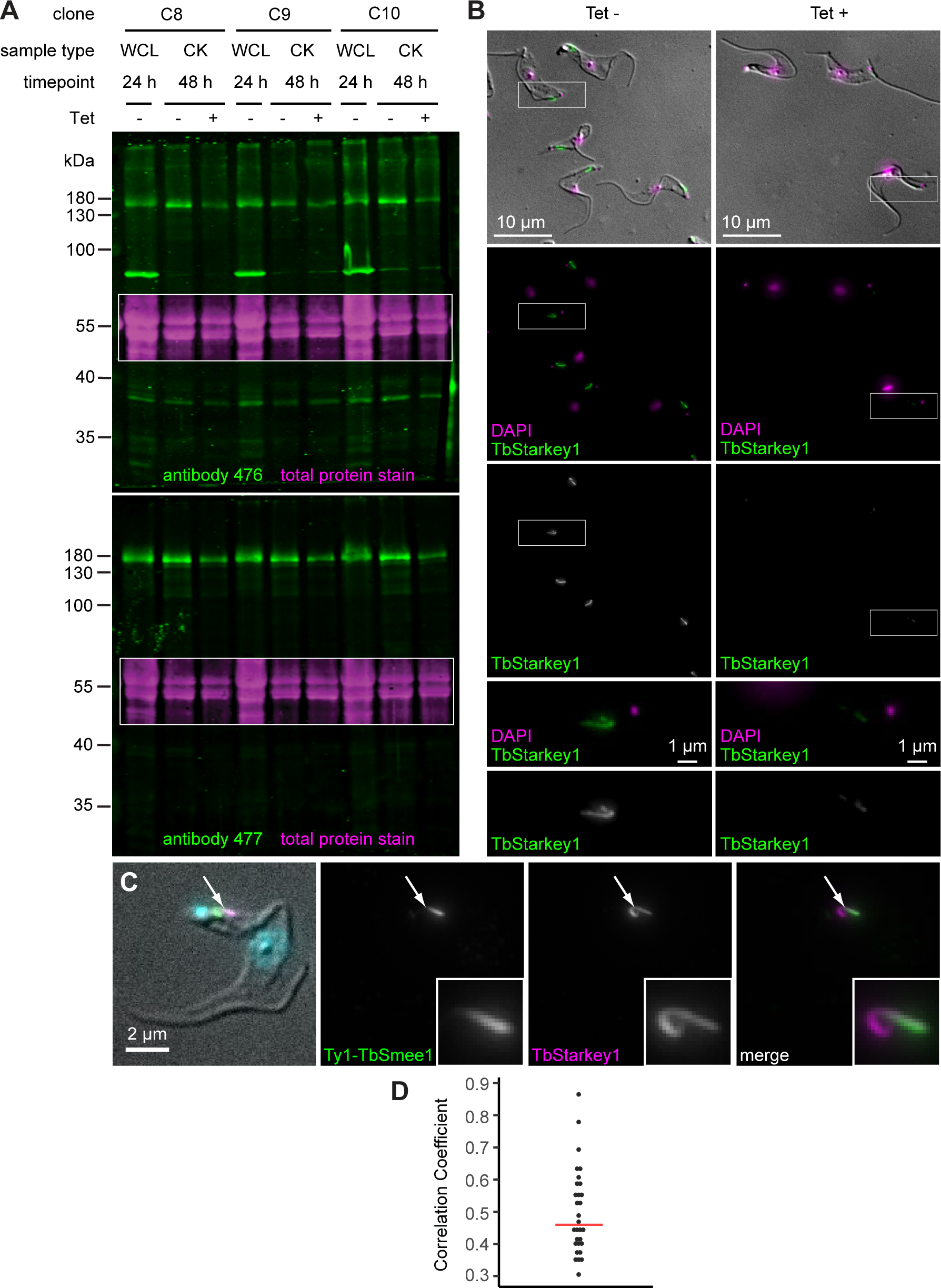
Validation of anti-TbStarkey1 antibodies. (A) Validation of anti-TbStarkey1 antibody specificity by immunoblotting. Three separate TbStarkey1 RNAi clones (C8, C9, C10) were analysed by immunoblotting. Both whole-cell lysates (WCL) and detergent-extracted cytoskeleton (CK) samples were obtained at the indicated timepoints (24 h, 48 h) from control (Tet-) and induced (Tet+) cells. The samples were immunoblotted using two separate anti-TbStarkey1 affinity-purified antibodies (antibody 476, 477). Both antibodies recognised a >130 kDa protein whose abundance was depleted after 48 h of RNAi. The 476 antibody additionally showed a significant cross-reaction with a protein of <100 kDa in WCL but not CK samples. A section of the total protein stain of each membrane is shown as an inset (magenta). (B) Validation of anti-TbStarkey1 antibody specificity by immunofluorescence microscopy. Control (Tet-) and TbStarkey1-depleted (Tet+) RNAi cells were extracted with non-ionic detergent, fixed with methanol, and labelled with anti-TbStarkey1 antibodies (green); DNA was stained using DAPI (magenta). TbStarkey1 localised to the hook complex, and signal was lost upon depletion. The boxed areas are shown enlarged in the bottom panels. Identical results were obtained using both anti-TbStarkey1 antibodies; exemplary images using the 477 antibodies are shown. (C) TbSmee1 overlaps with the shank part of the hook complex protein TbStarkey1 (arrow). All fluorescence images are maximum intensity projections, and an overlay with a single DIC section is shown. Overlap was manually confirmed in single z-slices. Results were obtained from multiple (n>3) independent experiments; exemplary images are shown. (D) Summary of measured correlation coefficients. Each dot represents a single cell; red lines show median values.

**Figure S5.**
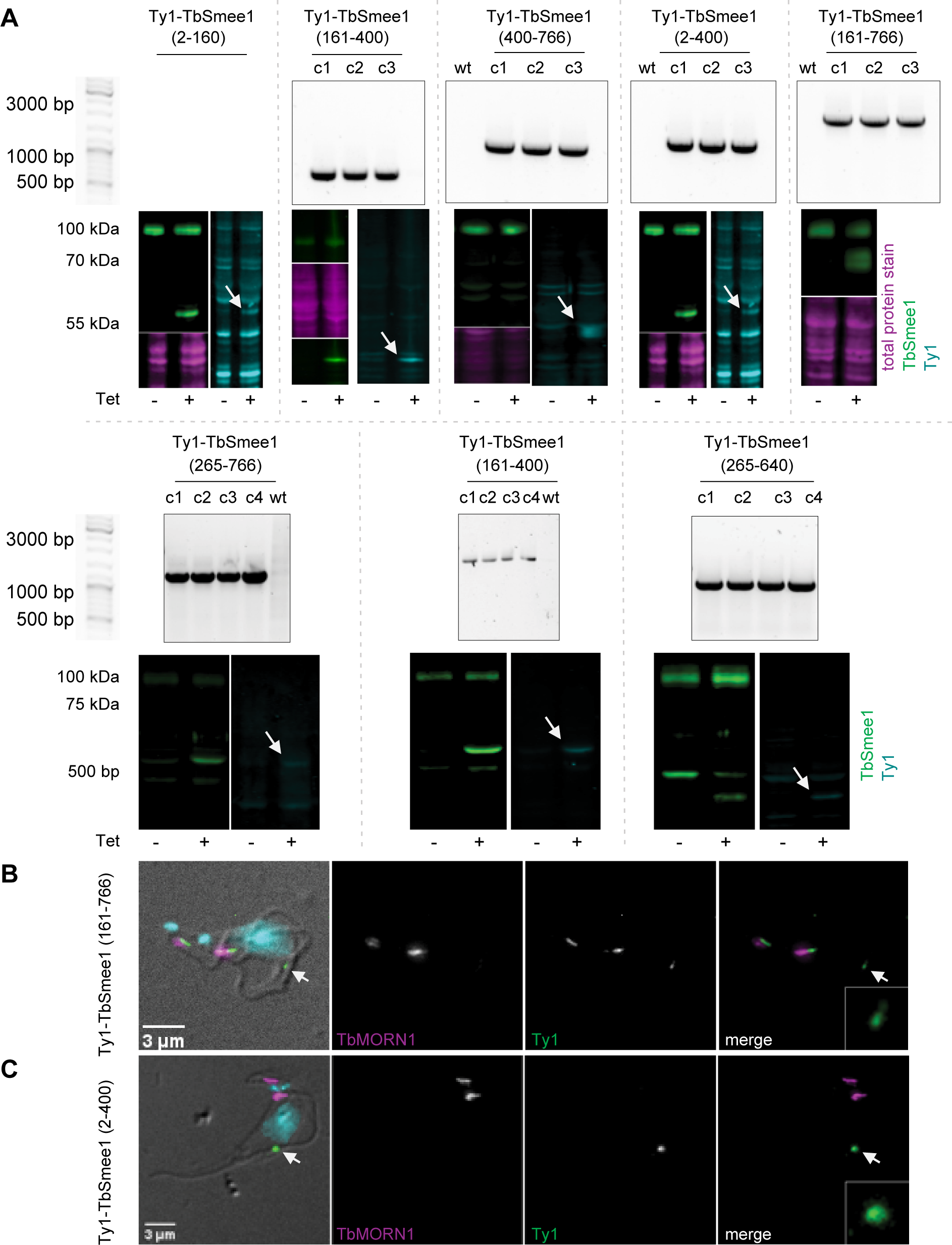
Validation of TbSmee1 truncations. (A) Validation of Ty1-TbSmee1 truncation cell lines by PCR analysis of genomic DNA and immunoblotting. The results for all 8 TbSmee1 truncations are shown, separated by dotted lines. Upper panels: PCR analysis of genomic DNA to confirm the presence of the truncation construct. PCR was used to amplify DNA from clones (c1, c2, c3) and wild-type control (wt) genomic DNA. Primers annealing to the sequence encoding Ty1 epitope and within the truncation were used. Lower panels: confirmation of Ty1-TbSmee1 truncation construct expression by immunoblotting. Whole-cell lysates from uninduced (−Tet) and induced (+Tet) cells were analysed by immunoblotting with anti-TbSmee1 (green) and anti-Ty1 (cyan) antibodies. Both antibodies detected proteins corresponding to the predicted size of the Ty1-TbSmee1 truncations; arrows indicate the target protein in the anti-Ty1 blots. A portion of the total protein staining of the membranes is shown in magenta. (B) Ty1-TbSmee1(161-766) localises to both the hook complex and the FAZ tip. Cells expressing the Ty1-TbSmee1(161-766) construct were extracted with non-ionic detergent, fixed with methanol, and labelled with anti-TbMORN1 and anti-Ty1 antibodies. DNA was stained with DAPI (cyan). Ty1-TbSmee1(161-766) was observed at both the hook complex and the FAZ tip (arrow). (C) Ty1-TbSmee1(2-400) localises to the FAZ tip but not the hook complex. Cells expressing the Ty1-TbSmee1(2-400) construct were extracted, fixed, and labelled as above. Ty1-TbSmee1(2-400) was observed exclusively at the FAZ tip (arrow). Images in panels B and C are maximum intensity z-projections with a single DIC z-slice overlay. Multiple (n>2) independent experiments using three separate clones for each construct were carried out; exemplary cells are shown.

**Figure S6.**
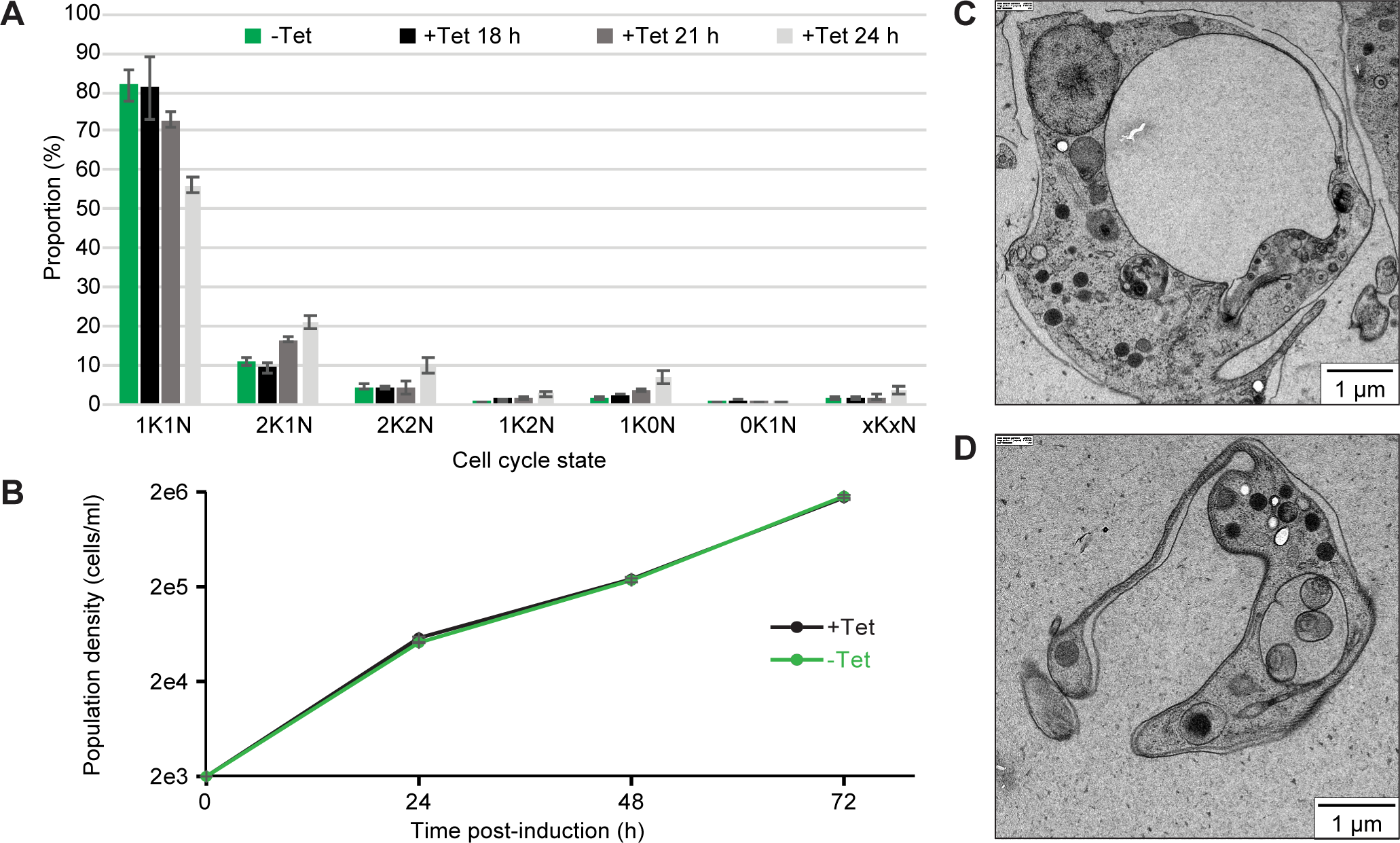
Additional results for TbSmee1 and TbStarkey1 RNAi experiments. (A) TbSmee1 depletion causes changes to cell cycle state distribution. Control (−Tet) and TbSmee1-depleted (+Tet) RNAi cells from 18 h, 21 h, 24 h post-induction were fixed using glutaraldehyde; DNA was stained using DAPI. The various cell cycle states (1K1N, etc) were manually quantified from images taken of the fixed cells. Data were obtained from three independent experiments, each using three separate clones; at least 240 cells were quantified for each timepoint. (B) Depletion of TbStarkey1 has no effect on population cell growth. Control (−Tet) and TbStarkey1-depleted (+Tet) cells were followed over a 72 h timecourse, and population density (cells/ml) was measured every 24 h. Data were obtained from three independent experiments, each using 3 separate clones. (C, D) Depletion of TbStarkey1 causes morphological abnormality. TbStarkey1-depleted cells were prepared for electron microscopy using high-pressure freezing and imaged. Cells with enlarged flagellar pockets could readily be observed.

**Figure S7.**
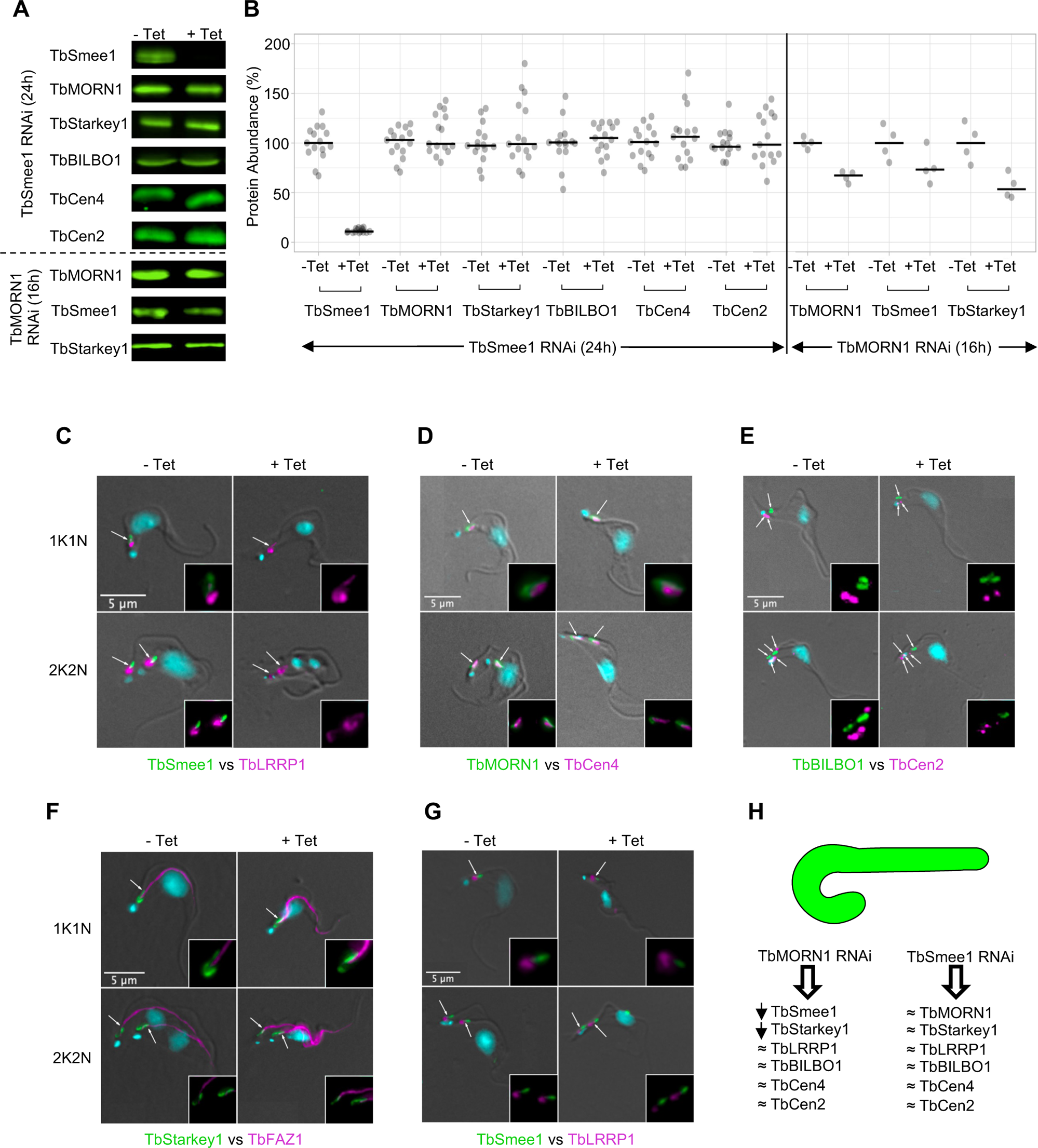
Depletion of TbSmee1 affects other cytoskeleton-associated proteins. TbSmee1 RNAi cells were induced for 24 h and analysed by immunoblotting of whole-cell lysates and immunofluorescence imaging of detergent-extracted cells. (A) Exemplary immunoblots showing the effect of 24 h TbSmee1 depletion on marker proteins for the hook complex, flagellar pocket collar, and centrin arm. For comparison, the effect of 16 h TbMORN1 depletion was also analysed. (B) Quantification of immunoblot data. TbSmee1 depletion did not affect any of the proteins analysed. Total protein staining was used for signal normalisation. Normalised TbSmee1 signals in +Tet cells were expressed relative to the −Tet signal for each sample. TbSmee1 −Tet signals were expressed relative to the mean of all TbSmee1 −Tet values in the dataset. Immunoblots from TbMORN1 RNAi experiments were quantified in the same way; TbMORN1 depletion resulted in a loss of both TbSmee1 and TbStarkey1 signal. The data shown were obtained from multiple (n>2) independent experiments, each using three (TbSmee1 RNAi) or two (TbMORN1 RNAi) separate clones. (C-F) TbSmee1 depletion does not affect the localisation of the marker proteins. Control (−Tet) and TbSmee1-depleted (+Tet) cells were extracted with detergent, fixed with methanol, and labelled with the indicated antibodies; DNA was stained using DAPI. The position of the hook complexes in exemplary 1K1N and 2K2N cells are shown with arrows; panel E also indicates basal bodies. Data were obtained from 2 independent experiments for each labelling combination, each using 3 separate clones. (G) TbSmee1 and TbLRRP1 in TbMORN1-depleted cells. (H) Summary of the observed effects (arrow = depletion, squiggles = no change) on the abundance of marker proteins caused by either TbMORN1 or TbSmee1 depletion, based on immunoblotting and immunofluorescence data.

**Figure S8.**
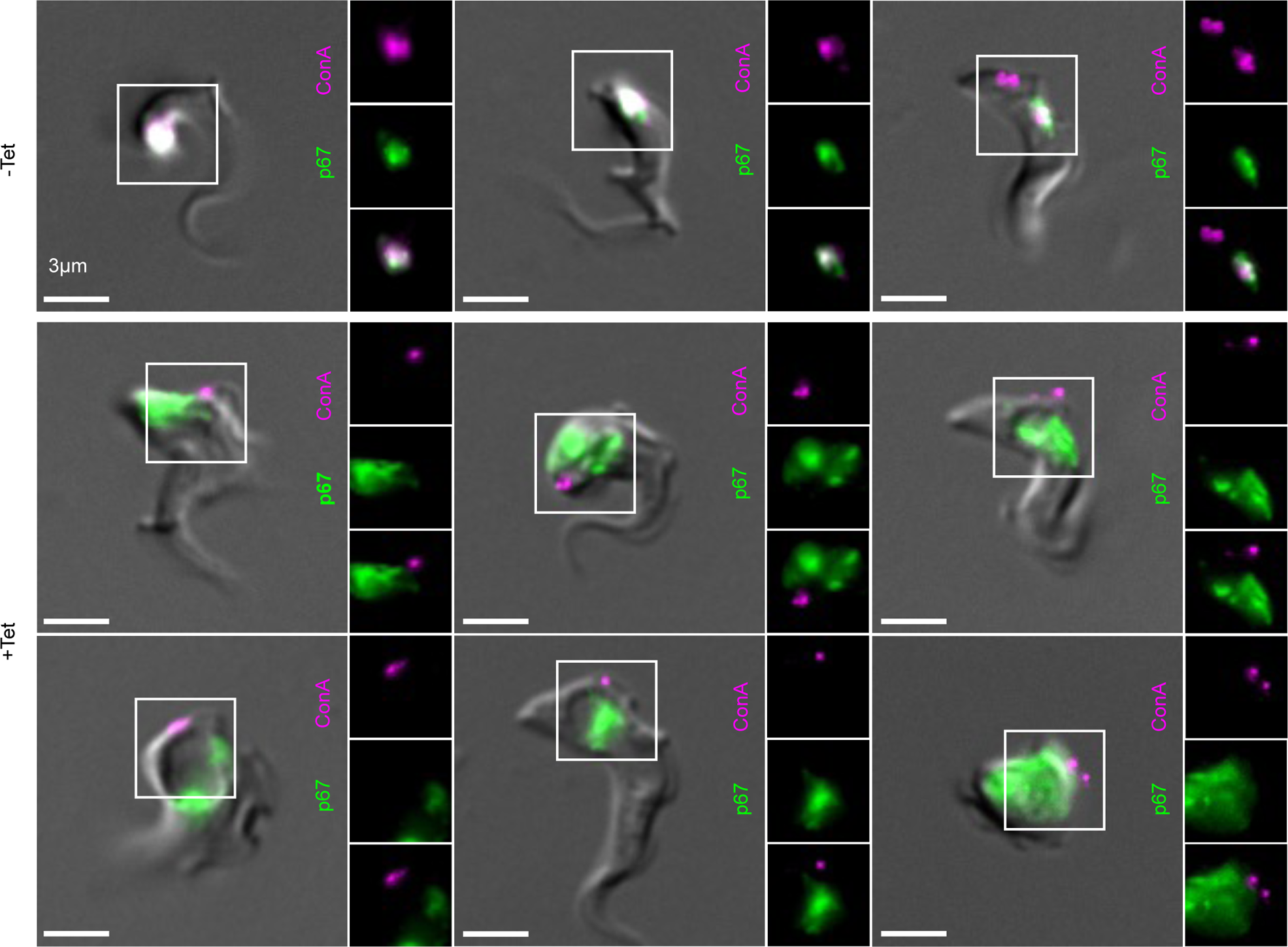
Depletion of TbSmee1 prevents trafficking of ConA to the lysosome. Control (−Tet) and TbSmee1-depleted cells (+Tet; 24 h timepoint) were incubated on ice with ConA (magenta). The cells were then shifted to 37 °C for 30 min to allow internalisation. The cells were then fixed and labelled with antibodies specific for the lysosome marker p67 (green). Control (−Tet) cells showed strong overlap between the two labels, indicating that ConA had been trafficked to the lysosome. +Tet cells showed no overlap between the two labels. Maximum intensity projections of the fluorescence channels are shown overlaid with a single DIC z-slice. Overlap in −Tet cells was confirmed in single z-slices. Single channels from the boxed area in each image are shown as insets. Data obtained from multiple (n>2) independent experiments each using three separate clones; exemplary cells are shown.

